# Genome-wide CRISPRa screens nominate modulators of CAR T cell survival within distinct tumor cytokine milieus

**DOI:** 10.1101/2024.03.16.583115

**Authors:** Benjamin C Curtis, Cailyn H Spurrell, Lindsay Flint, Aalton M Lande, Marissa Leonardi, James M Rosser, Ardizon Valdez, Nat Murren, Tiffanie Chai, Michael Fitzgerald, Jasmin Martinez-Reyes, Christopher P Saxby, Shannon K Oda, Michael CV Jensen

## Abstract

Chimeric Antigen Receptor (CAR) T cell therapy has revolutionized the treatment of B cell malignancies and translating this success to other cancers remains an ongoing clinical objective. Next-generation T cell products in development aim to genetically modulate many facets of cell behavior, for which gene-nominating platforms provide a useful framework for prioritization. Among competing screening approaches, CRISPR activation (CRISPRa) technology permits gain-of-function (GoF) gene surveys at genome-wide scale, but routine implementation in primary T cells has been stymied by high cell requirements (∼10^7^ - 10^8^) and abbreviated activity. Here, we describe a novel cell manufacturing schema using an all-in-one transposon-based gene delivery system coupled with CAR-restricted cell expansion to generate yields (10^9^) of primary T cells bearing CAR and CRISPRa transgenes that are well above the threshold needed for genome-scale screening. CRISPRa activity is sustained via the inclusion of divergent, duplicate Elongation Factor 1α core/human T-cell leukemia virus (EF1α-HTLV) hybrid promoters; while guide RNA representation is preserved through late lentiviral transduction, thus preventing bottlenecking and premature candidate pruning. CRISPRa-CAR T cells manufactured via this pipeline retain potent on-target gene-overexpression (>85% target^+^) across varied cell subsets (e.g. Tim-3^+^Lag3^+^ or serial-challenge) and timescales (>14 days). When deployed to survival-based genome-wide selection landscapes, CRISPRa-CAR pools nominate known and novel endogenous genes capable of enhancing CD8^+^ CAR T survival in cytokine-rich (e.g. *MYC, FUT6, IRF4, GSE1*) and cytokine-depleted (e.g. *CSF2RB*, *STAT6*, *IRF4*, *GSE1*) settings of tumor challenge. This system will have broad utility for therapy-enhancing gene discovery.

## INTRODUCTION

Chimeric antigen receptor (CAR) T cell therapy is an approved treatment for select hematological malignancies, but attempts to broaden its reach beyond cancers of B cell origin have produced limited success to-date.^1^ Consequently, emerging strategies for next generation therapy development often extend beyond refining CAR transgenes to include prospective genetic payloads that modify many different facets of cell behavior, with the ultimate aim of enhancing T cell potency.^2–6^ As gene candidates have accumulated, the burgeoning list of contenders has motivated efforts to streamline their evaluation using pooled molecular libraries, enabling researchers to test many prospective genes simultaneously.^7^

Despite their promise, gain-of-function (GoF) libraries composed of limited, pre-selected coding sequences (CDSs) restrict and implicitly bias the discovery process. Such collections are often weighted toward molecules with established roles in proximal T cell signaling or therapy resistance, and therefore exclude more distal or temporally-isolated genetic modifiers with the potential to steer treatment outcomes.^8^ In contrast, agnostic, discovery-based approaches to evaluating potency enhancement are slow, resource-intensive, and difficult to scale. While select research programs have sporadically surmounted these obstacles,^9^ molecular libraries composed of heterogeneous candidates suffer challenges of reproducibility and under-sampling, foremost due to variation in CDS length.^10^

To address these limitations, massively-parallel screening platforms based on CRISPR/Cas9 technology have emerged, operating at low cost and high scalability as a function of the system’s programmable, modular unit, the guide (g)RNA.^11^ This species can be readily packaged via lentiviral (LV) delivery platforms and delivered to primary T cells to modulate expression patterns of specific genomic loci.^12,13^ While traditional CRISPR-based screens concentrate on irreversible gene deactivation, this platform has been re-tooled to engage in broader gene modulation, which today includes transcriptional activation (CRISPRa), inhibition (CRISPRi), and various forms of epigenetic tuning.^14,15^

Applying the suite of updated CRISPR technologies to primary human cells, particularly lymphocytes, has been reportedly impeded by the inefficient gene delivery of the core homing component, the endonuclease-dead (d)Cas9.^12^ Despite this challenge, a lab recently demonstrated that CRISPRa^+^ antigen-agnostic primary T cells could be manufactured using high-quality lentivirus; when deployed to adaptive landscapes of limited duration, their platform nominated endogenous genes that potentiate therapeutically-relevant T cell functions.^16^ To realize their full potential however, such screens need to be applied to clinical T cell products and assessed over clinically-relevant timescales. Within these settings, CRISPRa holds tremendous promise as a gene-nomination platform to guide adoptive cell therapy product development. Here, we describe the development of a CRISPRa-enabling novel CAR T cell manufacturing method and subsequent first demonstration of a genome-scale GoF CAR T cell screens for therapy enhancement.

## RESULTS

Successful CRISPRa^+^ primary T cell manufacturing has been recently achieved using modified lentiviral delivery methods.^16,17^ Similar to most prior studies,^12^ our attempts with this approach yielded very low proportions of CRISPRa^+^ T cells and very low CRISPRa activity (**Fig S4**), despite deploying CDSs optimized for CRISPRa potency and positive cell identification (**Fig S1, Fig S2, Fig S3, supplemental results**).

Given our additional requirement to co-deliver a CD19CAR transgene, we considered if there were cell manufacturing schemas favoring simultaneous enrichment of CRISPRa and CAR transgene double-positive integrants. To this end, we adopted an all-in-one PiggyBac transposon-based delivery system (**Fig S5a**), utilizing donor plasmid DNA and mRNA coding for transient transposase. Recent studies have demonstrated that antigen-specific stimulation of transgenic T cells via (CD19) antigen^+^ cells from among peripheral blood mononuclear cell (PBMC) feeders can support incredible rates of cell-specific T cell expansion from rare founding populations.^18,19^

Following electroporation, gene-modified T cells were returned to a bed of autologous T cell-depleted PBMCs bathed in a cytokine-rich media, with selection initiated on Day 0 or 3 (**Fig 1a**). We term this cell manufacturing method “<u>E</u>lectro<u>p</u>oration-mediated <u>T</u>-cell Interface for <u>C</u>AR-limited <u>E</u>xpansion” (EP-TICLE). Via this process, we were able to generate approximately 100 and 10-fold expansion rates of CAR_gRNA_ and CRISPRa-CAR_gRNA_ T cells respectively, relative to bulk cell input (**Fig 1b, c**). We assume the true CAR-restricted expansion rates to be considerably higher. Notably, these specific-cell yields were >10-fold of those obtained via gene delivery following pan-T cell activation with stimulation beads.

**Figure 1:**
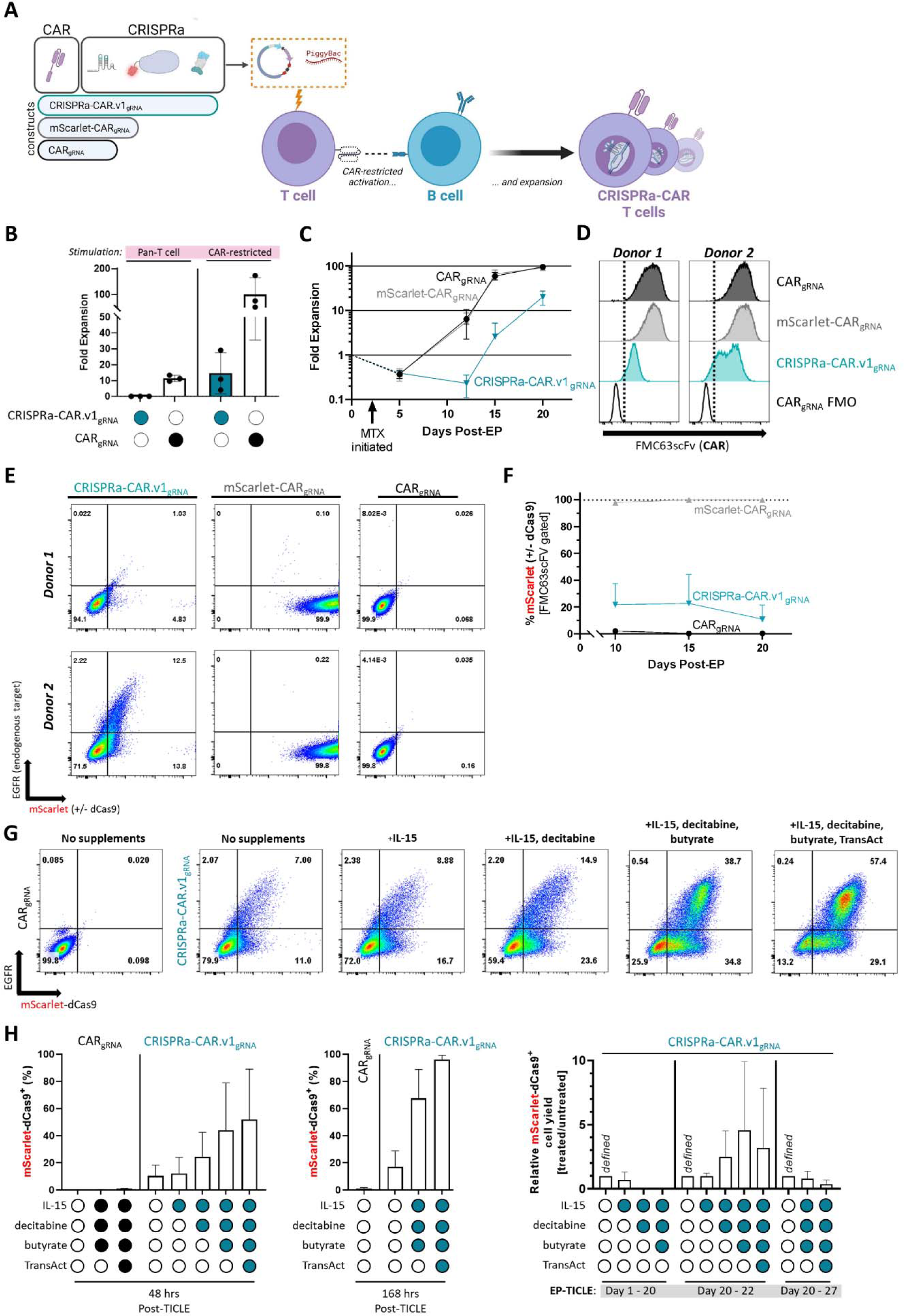
CRISPRa-CAR T cell EP-TICLE manufacturing and product characterization. (A) EP-TICLE manufacturing schema graphic. (B) Total live cell fold-expansion by Day 20 as a function donor plasmid and T cell stimulation method. (C) EP-TICLE live cell fold-expansion as function of time and donor plasmid. (D) Flow cytometry histograms depicting FMC63scFv (CAR) expression at Day 20 of EP-TICLE cell manufacturing as function of blood donor and donor plasmid. (E) Flow cytometry plots of EGFR (CRISPRa target gene) and mScarlet (+/−dCas9) expression in CAR, mScarlet-CAR, and CRISPRa-CAR.v1 CD8^+^ or CD4^+^ cells. (F) mScarlet positivity of EP-TICLE cells as a function of time and donor plasmid. (G) Flow cytometry plots of EGFR and mScarlet-dCas9 expression in CAR or CRISPRa-CAR.v1 mixed CD8^+^ and CD4^+^ T cells as a function of donor plasmid and 48-hour treatment with anti-silencing drug cocktails (e.g. IL-15, decitabine, butyrate, TransAct). (H) Quantification of mScarlet-dCas9 positivity or mScarlet-dCas9^+^ cell yield in CAR or CRISPRa-CAR.v1 as a function of donor plasmid, anti-silencing drug cocktail, and treatment time horizon indicated below each dataset. Data in (B), (C), (F), and (H) display the mean +/−SD of indicated measurement of cells generated from three human blood donors. Cells in (F) are gated on FMC63scFv^+^CD8^+^ or CD4^+^ live singlets. Cell plots or quantified data in (D), (E), (G), and (H) are gated on CD8^+^ or CD4^+^ live singlets only. All experiments depicted here used 1:1 mixtures of CD8^+^ and CD4^+^ T cells as EP-TICLE input.

### CAR-restricted T cell manufacturing can generate large numbers of CRISPRa-CAR T cells, but CRISPRa activity is limited

Intriguingly, despite achieving high CD19CAR purity in CRISPRa-CAR.v1_gRNA_ T cells (**Fig 1d**), expression of the CRISPRa transgene was persistently low, with poor induction of the CRISPRa target gene, the Epidermal Growth Factor Receptor (EGFR) (**Fig 1e, f, Fig S5b**). Subsequent relative vector copy number (VCN) analyses uncovered a modest reduction in CRISPRa_gDNA_ (18 - 29%) in 2 out of 3 donor cell products, although these frequencies could not account for the observed incidence of CRISPRa-negative cells (73 - 94%) as measured by flow cytometry (**Fig S5c - e**). Careful inspection of the donor plasmids failed to detect template impurities, suggesting that CRISPR ^−^ cells emerged *de novo* over the course of cell manufacturing, rather than as a result of contaminating, recombined DNA templates (**Fig S5g - i**).

We now considered if features of the CRISPRa CDS might be negatively impacting expression. For instance, foreign genes, regardless of their mechanism of delivery (e.g. lentivirus, transposon-based) often suffer from varying degrees of epigenetic silencing.^20,21^ By subjecting our CRISPRa-CAR.v1_gRNA_ cells to chemical species including IL-15; decitabine, a hypomethylating agent; butyrate, a histone deacetylase inhibitor; and TransAct™ (Miltenyi Biotec), a soluble T cell activation reagent, we observed additive increases in gene re-activation (**Fig 1g, h**, **Fig S6a, b**). Indeed, prolonged exposure (168 hrs) to anti-silencing drug cocktails led to near total restoration of CRISPRa positivity in CRISPRa-CAR.v1_gRNA_ donor cells (**Fig 1h**). Together, these results imply that dampened CRISPRa activity is primarily driven by severe DNA silencing of the CRISPRa transgene.

However, supplementation with epigenetically modifying drugs did not represent a viable path to rescue CRISPRa activity. For while short-term administration of these compounds led to marginal increases in specific cell yields, these trends turned net-negative after one week (**Fig 1h**). Similarly, continuous use of these reagents throughout primary manufacturing completely abolished effective cell expansion (**Fig 1h, Fig S6c, d**). Furthermore, since large doses of these drugs are not routinely prescribed concurrent with cell therapy, they would risk substantially confounding our screening results if introduced into subsequent selection landscapes.

Given the poor CRISPRa activity observed across multiple donor samples, we asked if CRISPRa toxicity might select for transgene loss. Prior studies for example have reported that high doses of either dCas9 or endogenous trans-activators have the potential to impose genotoxicity in various model systems, thus limiting cell viability.^22–24^

To probe for this suspected toxicity, we engineered a set of CRISPRa-deficient vectors to determine if transcription or subsequent translation of the CRISPRa transgene reduced T cell yields. Surprisingly, we observed no gross differences in relative cell counts derived from any CRISPRa construct variant (**Fig S7a, c**), suggesting that CRISPRa expression contributed no additive toxicity, and thus imposed no selective pressure favoring CRISPRa loss – a finding that aligns with other recent results.^25^ Finally, we tested the removal of the CRISPRa effector from the distal end of the CRISPRa transgene, but observed no increase in the proportion of mScarlet-dCas9^+^ cells, suggesting that the dCas9 CDS alone is sufficient to silence the entire transgene.

### Optimized CRISPRa transgene architecture resists deactivation

To sustain transgene expression, we now considered changes to our construct design, and evaluated various candidate promoters, cassette orientations, and re-positioning of our selectable marker (dihydrofolate reductase mutant, DHFR^FS^)(**Fig 2a**).^26^ Included in this survey were designs incorporating duplicate Elongation Factor 1α core/human T-cell leukemia virus (EF1α-HTLV) hybrid promoters, oriented divergently, rather than in tandem. To our surprise, this species dramatically increased CRISPRa expression (6-fold), purity (to 86%), and cell-specific yields (5-fold) (**Fig 2b**). No equivalent increase in performance was observed with alternative promoter combinations or orientations (**Fig S8a - d**). CRISPRa expression was further boosted by transferring the selectable marker from the CD19CAR-containing coding cassette to the N- or C-terminus of the CRISPRa CDS, with the C-terminal DHFR^FS^ positioning yielding slightly higher CRISPRa positivity (97%) and increased intensity (+2.4-fold) without sacrificing cell yield (**Fig 2b**).

**Figure 2:**
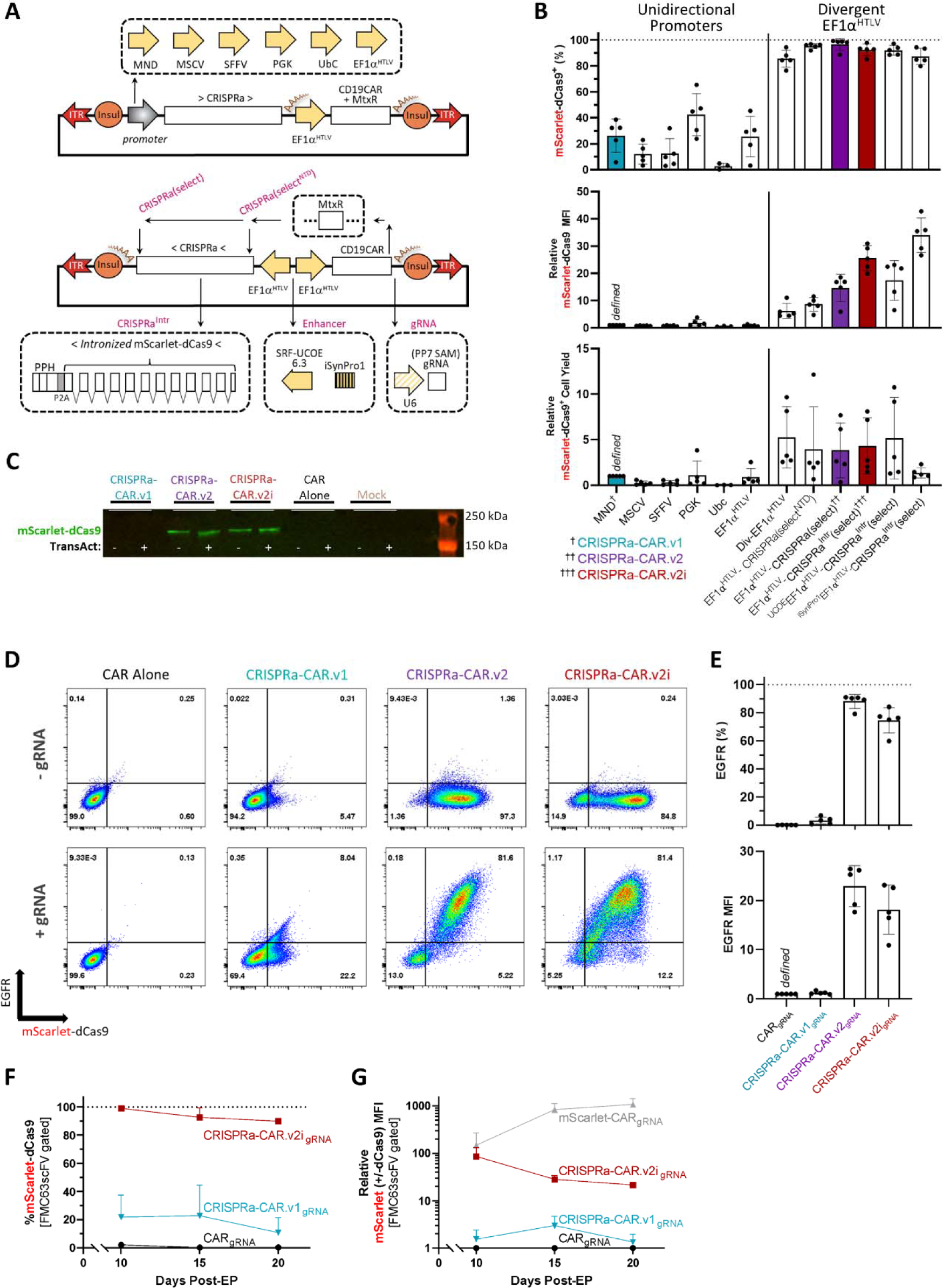
Optimization of CRISPRa-CAR vectors. (**A**) Abridged donor plasmid maps of select CRISPRa-CAR variants. (**B**) Quantification of Day 20/21 mScarlet-dCas9 positivity (top graph), relative mScarlet-dCas9 mean fluorescence intensity (middle graph), and relative mScarlet-dCas9^+^ T cell yield (bottom graph) of indicated CAR T cells generated via EP-TICLE. (**C**) mScarlet-dCas9 immunoblot of untransposed mock, CAR, CRISPRa-CAR.v1, CRISPRa.v2, and CRISPRa-CAR.v2i EP-TICLE cells at Day 20/21. (**D**) Day 20 flow cytometry plots of EGFR and mScarlet-dCas9 expression in cells generated via EP-TICLE from indicated donor plasmid with or without accompanying all-in-one U6-gRNA(EGFRprom) expression cassette. (**E**) Day 20/21 quantitative EGFR positivity (top graph) and EGFR mean fluorescence intensity (bottom graph) of cells generated via EP-TICLE from indicated donor plasmid. (**F**) mScarlet-dCas9 positivity and (**G**) relative mScarlet(+/−dCas9)^+^ cell yield of CAR, mScarlet-CAR, CRISPRa-CAR.v1_gRNA_, and CRISPRa-CAR.v2i_gRNA_ T cells as a function of time and donor plasmid. Cells in (B), (D) and (E) are gated on CD8^+^ or CD4^+^ live singlets only. Cells in (F) and (G) are gated on FMC63scFv^+^CD8^+^ or CD4^+^ live singlets. Data shown in (F) for CRISPRa-CAR.v1_gRNA_ cells is copied from Fig 1c and has not been generated from a distinct experiment. All experiments depicted here used 1:1 mixtures of CD8^+^ or CD4^+^ T cells as EP-TICLE input. Quantified data in (B) and (E) is compiled from cells generated from five human blood donors. Time course data in (F) and (G) display the mean +SD of indicated measurement of cells generated from three human blood donors.

Drawing from recent reports highlighting the success of including scattered introns throughout the Cas9 transgene to boost or sustain expression levels,^20,27,28^ we tested installing 11 short human introns within the mScarlet-dCas9 cassette, drawn from essential housekeeping, cell cycle, tumor suppressor, or T cell-enriched genes. Cells manufactured using this candidate vector expressed greater dCas9 levels (+1.8-fold) with only a slight reduction in overall positivity (92%) relative to the non-intronized equivalent. Lastly, we tested the deployment of two enhancer-like elements: a ubiquitous chromatin opening element (SRF-UCOE6.3)^29^ or a synthetic response element (“iSynPro1”) engaged by CAR-ligation (**Fig 2b, Fig S9a - c**).^30^

Based on our findings, we selected two final candidates for further characterization: a CRISPRa construct with divergent EF1α-HTLV promoters and DHFR^FS^ in-frame with the CRISPRa polycistron (“CRISPRa-CAR.v2”), and the intronized version of the same (“CRISPRa.CAR.v2i”). In contrast to CRISPRa-CAR.v1_gRNA_, cells generated from either of these optimized candidate vectors evidenced much higher activity as judged by EGFR positivity and MFI (**Fig 2d, e, Fig S9d**).

CRISPRa-CAR.v2i cells maintained high levels of CRISPRa expression throughout EP-TICLE manufacturing without transgene loss (**Fig 2c, f, g, Fig S5c, d**). CRISPRa-CAR.v2i cells were also successfully generated from CD4^+^ or CD8^+^ monocultures and achieved both similar cell yields and CRISPRa target expression levels compared to EP-TICLE cultures seeded with 50:50 mixtures of CD4^+^/CD8^+^ T cells (**Fig S9d**). In sum, our second-generation transposon-based vectors could generate potent CD4^+^ or CD8^+^ CRISPRa-CAR T cells at sizable quantities (> 10^9^) well in excess of the requirements for genome-scale screening (≈ 10^7^ - 10^8^).

### gRNA can be delivered to CRISPRa-CAR T cells ***in trans*** following primary manufacturing

Given the extended duration of cell manufacturing (∼ 3 weeks) and magnitude of cell-specific expansion (estimated to be >100-fold), we were concerned about the potential for significant and unpredictable distortion of gRNA representation. Specifically, culture bottlenecking and subsequent drift might skew the distribution of gRNA^+^ clones, while premature CRISPRa activity would unintentionally shape and prune gRNA frequencies early in cell manufacturing. To address both concerns, we investigated introducing gRNAs late in primary culture. To execute this strategy, we once again turned to lentivirus.

Conventionally, CAR T cells are manufactured by lentiviral transduction shortly following T cell activation; stimulated cells are primed for successful viral integration due in part to their higher metabolite availability,^31^ dearth of restriction factors,^32^ and upregulation of the low-density lipoprotein receptor.^33^ We hypothesized that T cells manufactured via EP-TICLE might retain sensitivity to lentiviral transduction despite their chronological distance from stimulation, as a result of their sustained proliferation.

Nevertheless, as lentivirus transduces quiescent T cells at vastly diminished efficiencies relative to their recently-activated counterparts,^34–36^ we also evaluated pseudotyped lentiviral delivery systems featuring viral envelopes derived from Measles (H/F) and Baboon endogenous retrovirus (BaEVRless) (**Fig 3a**). These variants possess high affinity for cell receptors decorating the surface of resting T cells.^37,38^ In parallel, we also tested a modified envelope featuring an anti-CD3scFv, capable of coupled T cell stimulation and transduction (**Fig 3a**).^39^

**Figure 3:**
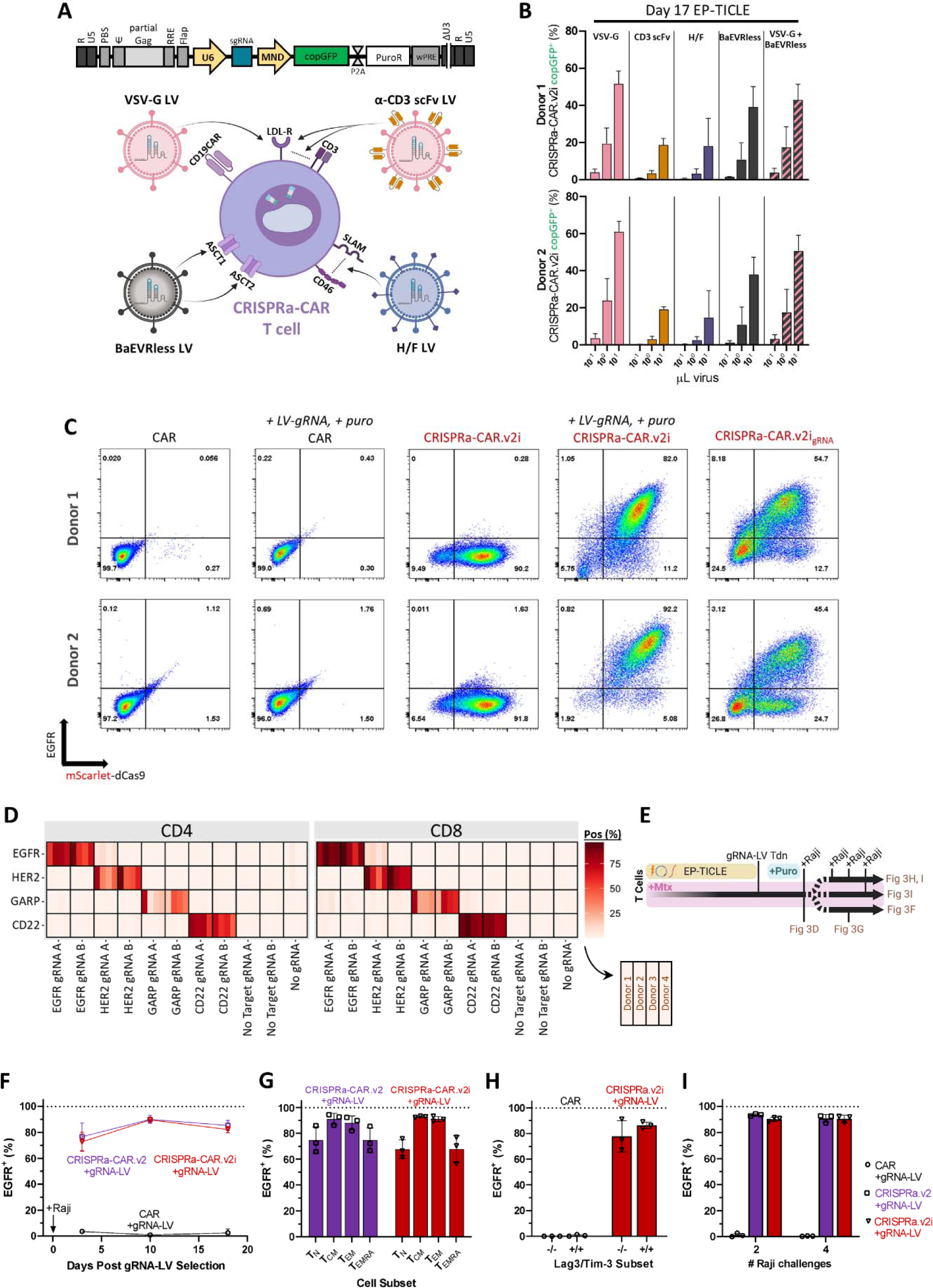
gRNA-LV transduction of CRISPRa-CAR T cells. (A) Abridged map of ssRNA lentiviral genome (*top*) and graphic depicting pseudotyped lentiviral virions with envelope cognate receptor on CRISPRa-CAR T cells (*bottom*). (B) copGFP (gRNA) positivity of CRISPRa-CAR.v2i T cells as a function of viral packaging using designated viral envelope variant (*top*) and volume of concentrated lentivirus. Transductions were performed on Day 17 or 21 of EP-TICLE (indicated) using cells manufactured from two blood donors. (C) Flow cytometry plots of EGFR and Scarlet-dCas9 expression in cells generated via EP-TICLE and where indicated, puromycin-selected after transduction with (U6p-)gRNA-lentivirus. (D) heat map of indicated gRNA target gene protein expression as a function of gRNA-LV across CD4 (left) and CD8 (right) in CRISPRa-CAR.v2i T cell subsets and among four separate human donor sourced cells generated via EP-TICLE and distal lentiviral transduction. gRNA “A” and B” contain distinct protospacer sequences specific to each respective gene’s promoter. (E) Cell manufacturing and experimental schema for data displayed in figures (F) thru (I). (F) Quantification of EGFR positivity in CAR-CRISPRa or CAR T cells after EP-TICLE and subsequent distal transduction. (G) Proportion of EGFR^+^ CRISPRa-CAR T cells in designated differentiation class (T_N_, CD45RO^−^CD62L^+^; T_CM_, CD45RO^+^CD62L^+^; T_EM_, CD45RO^+^CD62L^−^; T, CD45RO^−^CD62L^−^) after manufacturing and target cell stimulation. (H) Proportion of EGFR^+^ T cells as a function of inhibitory receptor positivity (Lag3, Tim-3) and donor plasmid after cell manufacturing and serial tumor challenge.) (I) Proportion of EGFR^+^ CRISPRa-CAR or CAR T cells as a function of number of CAR target cell line challenges after EP-TICLE and subsequent distal transduction. Cells in (D) were selected with puromycin (except “no gRNA”) for 48 hours and were then co-incubated with Raji cells for 48 hrs at a 1:2 effector-to-target ratio. Cells in (C) are gated on CD8^+^ or CD4^+^ live singlets. Cells in (D) are gated on mScarlet-dCas9^+^CD8^+^ or CD4^+^ live singlets. Cells in (F) and (I) are gated on copGFP^+^CD8^+^ or CD4^+^ live singlets. Cells in (G) are gated on copGFP^+^mScarlet-dCas9^+^ CD8^+^ or CD4^+^ live singlets. Cells in (H) are gated on CD8^+^ live singlets. Plotted values in (F) thru (I) are the mean +/−SD of cells from three human donors. In plots (F) thru (I) the following cell groups are designated by the indicated shape: CAR (), CAR + gRNA (), CRISPRa-CAR.v2 + gRNA (), CRISPRa-CAR.v2i + gRNA ().

To our surprise, CD19CAR T cells intercepted starting on Day 17 of EP-TICLE transduced most efficiently on a per volume basis with lentivirus packaged with the conventional vesicular stomatitis virus glycoprotein (VSV-G), although BaEVRless-pseudotyped LV transduced cells at similar rates (**Fig 3b, Fig S10g - h**). Subsequent optimization of T cell plating conditions increased late transduction efficiencies of EP-TICLE cells further still (**Fig S11b - f**). In a complementary fashion, optimization of the gRNA transfer vector also boosted gRNA-LV titer and overall CRISPRa activity levels (**Fig S10a - f, Supplemental results**).

To pair this new delivery vehicle with our manufacturing process, CRISPRa-CAR T cells were generated via EP-TICLE and then transduced with gRNA-LV (Libr.1) around Day 17. Cells produced in this fashion were shown to exhibit similar levels of target gene activation compared to cells receiving an *cis* gRNA without secondary transgene delivery (**Fig 3c**). Leveraging this novel combination manufacturing schema of early EP and late LV transduction, cells were made to express a variety of target gene products normally absent on conventional T cells. High, specific, and reproducible overexpression was confirmed, with modestly higher activity levels seen in CD8^+^ T cells (**Fig 3d, Fig S11a**).

### CRISPRa-CAR**_gRNA_** T cells exhibit sustained CRISPRa activity across various timescales and cellular phenotypes

To determine possible selection landscapes compatible with our CRISPRa-CAR_gRNA_ T cells, we examined the frequency and duration of CRISPRa activity within T cell subsets of clinical interest (**Fig 3e**). Following cell manufacturing, both CRISPRa-CAR.v2_gRNA_ and CRISPRa-CAR.v2i_gRNA_ T cells evidenced robust and stable on-target gene expression (EGFR) over an additional two weeks of culture, with only a minor loss of potency upon methotrexate (Mtx) removal (**Fig 3f, Fig S12a**). All T cell differentiation subsets exhibited robust CRISPRa target overexpression, although central (CD45RO^+^CD62L^+^) and effector memory (CD45RO^+^CD62L^−^) subsets displayed modestly higher CRISPRa target activity than naïve-like (CD45RO^−^CD62L^+^) or terminal effector (CD45RO^−^ CD62L^−^) subsets (**Fig 3g, Fig S12b**). In contrast, T cells serially challenged with CAR targets or bearing high levels of immunoinhibitory receptors (Lag3^+^Tim-3^+^) exhibited no differences in CRISPRa activity compared to cells subjected to fewer challenges or lacking these phenotypic markers (Lag3^−^Tim-3^−^), respectively (**Fig 3h, i, Fig S12c, d**).

### Genome-wide CRISPRa screens identify unknown modulators of CAR T cell persistence

To demonstrate the clinical utility of our novel gene-nomination platform, we chose to measure CAR T cell persistence following assignment to one of three distinct environments: (1) T cell monoculture, (2) Raji challenge with cytokines, and (3) Raji challenge absent cytokines. To implement, CRISPRa-CAR CD8^+^ T cells were manufactured via EP-TICLE and late in culture, underwent lentiviral transduction using subcloned CRISPRa gRNA-LV half-libraries (**see methods**). Following secondary selection, cells were assigned to treatment arms, cultured for seven days, and collected for processing (**Fig 4a, f, Fig S13b - e**).

**Figure 4:**
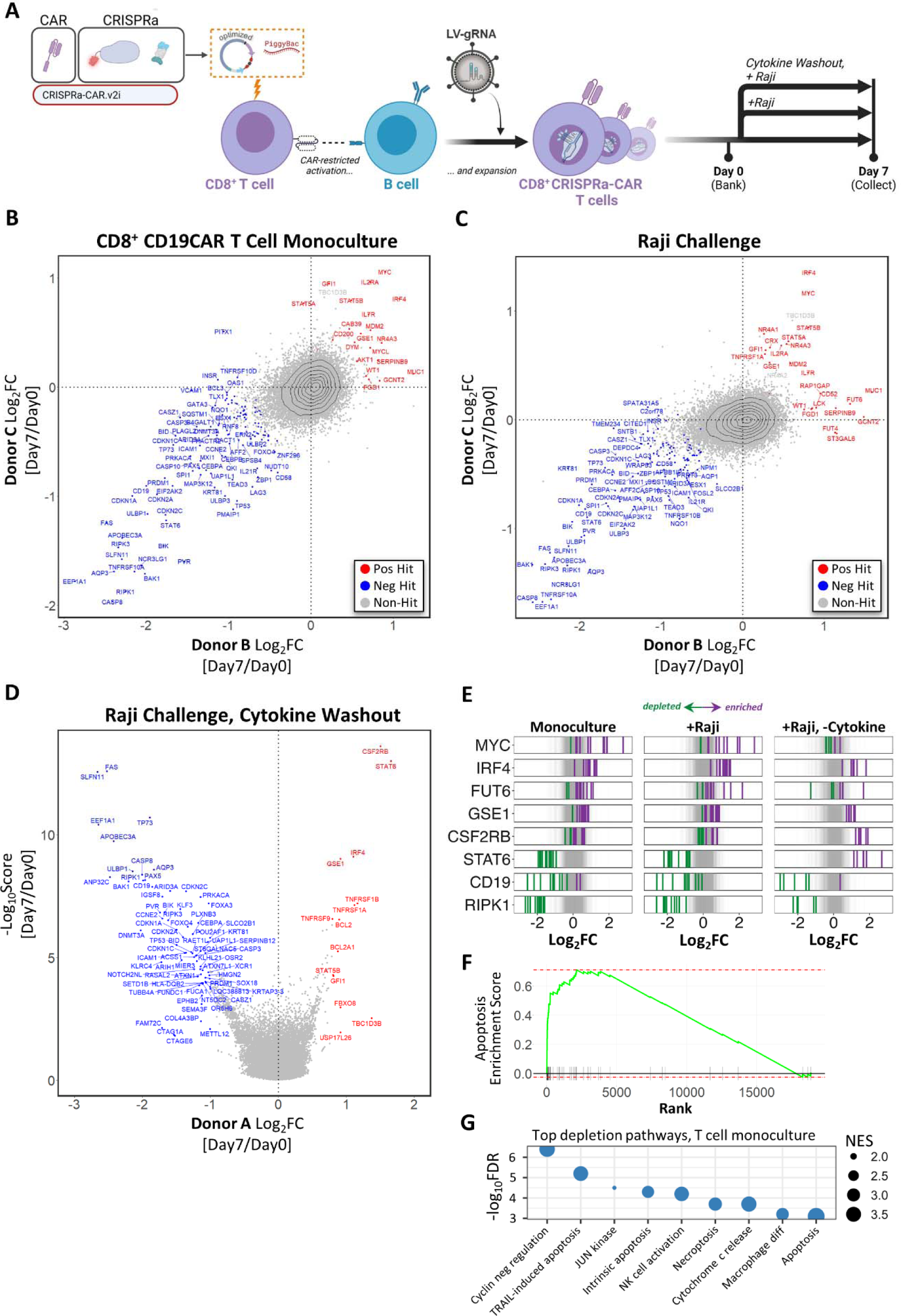
CRISPRa-CAR T cells genome-wide persistence screen results. (**A**) Schematic of genome-scale CRISPRa-CAR T manufacturing and landscape treatment arms. (**B, C**) Plot of gRNA median Log_2_fold-change (FC) of Donor B and C from Day 0 to Day 7 for the (B) cytokine-supplemented monoculture (C) and cytokine-supplemented T/tumor cell co-culture arms. (**D**) Volcano plot of gRNA median Log_2_FC (x-axis) vs. −Log_10_Score (y-axis) of Donor A from Day 0 to Day 7 for the T/tumor co-culture with cytokine washout. (**E**) Distribution of individual gRNAs across treatment arms for specified gRNA-targeting gene sets. (**F**) Gene set enrichment analysis for the apoptosis pathway, ordered from most- to least-depleted gene-specific gRNA set. (**G**) Graphic depicting top-ranking non-redundant gene pathway categories where gene depletion is most concentrated at indicated normalized enrichment score (NES). Positive (red) and negative (red) hits in (B, C, D) indicate statistically significant sets of gene-specific gRNAs with FDR < 0.05 (B, C) or adjusted p value < 0.05 (D). Purple and green gRNAs in (E) indicate enriched or depleted gRNAs, respectively. Gray shading in (E) indicates distribution of non-targeting gRNAs for each treatment arm. Analyses for (F) and (G) are drawn from the T cell monoculture conditions only.

Following cell recovery, genomic (g)DNA was extracted, and gRNA protospacer sequences were PCR-amplified and then tabulated via next generation sequencing (NGS) (**Fig S14a**). Following quality assessments (**Fig S14b - d**), Day 0 samples were compared to Day 7 samples across all three treatment arms, using the MAGeCK algorithm.^40^ From this analysis emerged a collection of differentially accumulated (“positive hits”) or diminished (“negative hits”) gene-specific gRNAs as measured by median log_2_-fold change (FC) (**Fig 4b - d**).

As expected, gRNAs targeting genes known to potentiate cell death pathways were highly diminished at Day 7 (**Fig 4f, g**). Negative hits included known regulators of cell intrinsic death (e.g. *BID*, *BIK*, *BAK1*), cell extrinsic death (e.g. *FAS*, *TNFRSF10A*, *CASP8*, *CASP10*), common cell death pathway (e.g. CASP3), and cell cycle inhibitors (e.g. *CDKN1A*, *CDKN1C*, *CDKN2A*, *CDKN2C*, *TP53*, *TP73*) (**Fig 4b - d**). Unlike for prior CRISPRa dropouts screens, here *CD19* arose as a strong negative hit, and suggests both CRISPRa and CD19CAR functionality via “programmed fratricide.” Similarly, *EEF1A1* (*EF1α*), also arose as an unusual negative hit across treatment arms, which we hypothesize to be an artifact of our optimized transposon-based payload, with the CRISPRa system engaging with the integrated CRISPRa-CAR.v2i duplicate EF1α-HTLV hybrid promoters, driving a lethal positive feedback loop. In total, the large constellation of depleted gene-specific gRNAs suggests that our CRISPRa screen achieved gene over-expression levels sufficient to markedly modulate cell phenotypes.

As positive hits between the T cell monoculture and cytokine-supplemented Raji challenge treatment arms were extensively overlapping (**Fig S14c**), we summarize them first together (**Fig 4b, c**). Tellingly, positive hits included multiple known proto-oncogenes (e.g. *MDM2*, *MUC1*, *MYCL*, *WT1*, GF1),^41–46^ including the two canonical members of the *IRF4*-*MYC* pathway.^47–50^ Also represented were two gamma chain cytokine receptor subunits (e.g. *IL2RA*, *IL7R*) and their downstream mediators (e.g. *STAT5A*, *STAT5B*, *AKT1*). Other nominees included *SERPINB9* and *TNFRSF1A* (TNFR1), which have been shown to protect T cells from distinct mechanisms of activation-induced cell death.^51,52^ A surprising collection of positive hits across cytokine-supplemented treatment arms included members of the NR4A orphan nuclear receptor family of transcription factors (e.g. *NR4A1/Nur77*, *NR4A3/NOR1*); under canonical models, these early T cell response genes serve to dampen T cell proliferation following activation.^53–55^ Other positive hits associated with T cell activation included *JUNB*, *LCK*, *CD52*, and *CD200*.^56–60^ Importantly, also enriched were positive hits without defined roles in T cell survival (e.g. *GSE1*, *GCNT2*, *DYM*, *FGD1*, *CAB39*, *CRX*, *RAP1GAP*), suggesting that our screening platform is capable of nominating genes with unrecognized therapeutic potential. Lastly, hits unique to cytokine-supplemented Raji co-culture include transferases known to modulate leukocyte cell-to-cell adhesion (e.g. *FUT4*, *FUT6*, and *ST3GAL6*).^61,62^

The results of the Raji challenge without cytokine nominated positive hits that were both overlapping (e.g. *IRF4*, *TNFRSF1A*, *GIF1*, *STAT5B*, *GSE1*) and distinct (e.g. *CSF2RB*, *STAT6*, *TNFRSF1B*, *TNFRSF9*, *BCL2, BCL2A, FBXO8, USP17L26*) from those found under cytokine-rich conditions (**Fig 4d**). One of the strongest positive hits, *CSF2RB*, encodes the cytokine receptor common subunit beta, CSF2RB; an activating mutation of this gene have been reported to drive cytokine independent growth in a patient case of T cell lymphoma.^63^ Other positive hits include *TNFRSF1A* and *TNFRSF1B* (TNFR2), the products of which together which form an antiapoptotic heterocomplex, and both having been nominated by the one prior human T cell genome-scale CRISPRa screen.^16^ Finally, *STAT6* paradoxically registered as either a positive or negative hit within the cytokine-free and two cytokine-rich arms, respectively. Exogenous IL-4 was a component part of our cytokine-supplemented media, and has been shown to coordinate with STAT6 signaling, directing downstream nutrient metabolism;^64–67^ therefore, we suspect that IL-4 and/or another exogenous cytokine (i.e. IL-2, IL-7, IL-21) dictates whether *STAT6* overexpression favors CAR T cell death or survival.

In total, these results demonstrate the capacity of our screening platform to nominate endogenous genes that enhance CAR T cell fitness in distinct environments.

## DISCUSSION

### A new tool to guide therapy enhancement

In the last decade, CAR T cell therapy has advanced considerably from its underperforming first-generation aspirants^68^ to today comprising multiple clinical treatment options.^69^ And yet, T cell therapies require further development if they are to provide predictable, durable remissions for the most prevalent forms of cancers. In this study, we describe a novel CRISPRa-CAR T cell manufacturing method, enabling longitudinal screens and ultimately nominating novel mediators of CAR T cell survival in the presence of tumor and variable cytokine levels.

Of note, while a recently published CRISPRa screen of human primary T cells identified a highly similar set of pro-apoptotic (negative hit) genes,^16^ our list of pro-survival (positive hit) genes is unique to our specific screening platform. We suspect that this may be partially a feature of benchmarking our analysis to a Day 0 sample following cell manufacturing, rather than to the gRNA library plasmid pools. In the latter scenario, we speculate that measured T cell persistence is more a readout of cell product manufacturing fitness. To our knowledge, this is the first genome-scale over-expression screen undertaken in CAR T cells, or any alternative clinically-oriented primary T cell product.

### Promoter organization and silencing resistance

One of our key findings was the observation that divergent dual EF1α-HTLV promoters markedly protected their transgene dependents from silencing, possibly mediated by the HUSH silencing complex.^20^ This result aligns with research demonstrating that native divergent promoters may better resist heterochromatin formation.^70,71^ During our exploration of further divergent promoter pairings, only EF1α-HTLV/MND (“Div-MNDrvs”) with EF1α-HTLV aimed at the CRISPRa transgene behaved similarly (**Fig S8d**). However, this also suggests that both promoters need not be identical in future designs to extract the same benefit, so long as EF1α-HTLV drives transcription of the susceptible CDS.

Although gross donor plasmid impurities were not seen in templates used for cell manufacturing (**Fig S5f - i**), we did witness non-trivial rates of recombination following routine bacterial transformation using our nanoplasmid-competent cell line. As a result, we cannot exclude the remote possibility that trace amounts of recombined plasmid, undetected in our preparations, may contribute to the sporadic CRISPRa transgene loss observed in a portion of select donor cells.

### dCas9 transgene silencing limits CRISPR utility

Given the evidence of transgene silencing occurring following transposon-based delivery, we questioned whether our early lentiviral attempts had been similarly affected. VCN analyses of dual-transduced (dCas9 LV + CAR-PPH_gRNA_ LV) T cells revealed mean CRISPRa-to-CAR transgene ratios of 1.2 (**Fig S6f**). Perhaps unsurprisingly then, subsequent treatment of these same cells late in culture with butyrate and TransAct™ drove mScarlet-dCas9 levels to as high as 64% positivity (**Fig S6e**). Together, these results hint that poor dCas9 expression reported in numerous studies, having been attributed to mediocre dCas9 lentiviral delivery, may more generally be affected by the silencing of dCas9 transgenes. Given that HUSH templates include episomal as well as integrated DNA,^20^ this phenomenon may thus also hinder vRNA packaging by producer cells, and thereby would favor dCas9-lentiviral production timelines with accelerated supernatant harvesting.^16^

Furthermore, CRISPRa/i-based adaptive landscapes that limit culture duration or activate T cells as an obligate screening component may mask underlying challenges in long-term dCas9 expression. The designs presented here will enable more complex, longitudinal screens without the added need for extensive donor pre-screening. As such, we expect our gene-nomination platform to more closely mirror the likely activity of translated gene cassettes, and not be biased towards a subset of patients or cells with overly-permissive expression of foreign DNA.

Recently, labs have sidestepped these difficulties in sustaining CRISPRa activity, by instead delivering an alternative minimal dCas9 ortholog derived from *Staphylococcus aureus*.^17^ At present, it is unclear how such methods compare to our own in terms of activity level and duration.

More generally, our challenge of reinforcing dCas9 expression likely represents a specific instance of broad vulnerability in transgene therapy. Specifically, silencing of foreign DNA is a non-trivial problem, occurring in primary T cells at high frequency.^72^ And yet, arrayed and pooled screening systems routinely deploy CDSs with uncharacterized long-term durability.^9^ For this reason, pooled, constitutive open reading frame (ORF) screens are expected to incur non-negligible numbers of false negatives as a function of increasing CDS size – a candidate type already underrepresented (and therefore underpowered) in most ORF libraries, even at the moment of landscape initiation.

### CRISPRa-CAR T cell… therapy?

To reiterate, the primary aim of our platform development has been to enable the discovery of novel endogenous regulators of therapeutic relevance, and then translate these native genes to common ORFs or rationally-designed synthetic genes, intended for CAR or other forms of adoptive T cell therapy. However, given their modularity and capacity for multiplex gene modulation,^73^ we envision that CRISPRa^+^ T cells could conceivably be offered, either as stand-alone or programmed, personalized GoF treatment options in due course. Indeed, private pharmaceutical companies have been founded, predicted on the development of *in vivo* CRISPRa drugs. Given the hints of preexisting anti-Cas9 humoral and cellular immunity within the human population,^74–76^ CRISPRa commercial drug development has thus far been restricted to immune-privileged anatomical sites, such as the central nervous system.^77^

### Future CRISPR screening in T cells and beyond

Survival-based CRISPRa readouts were permissible here without cell sorting due to the low frequency of gRNA^+^ CRISPRa-negative cells (**Fig S13d**), the presence of which would dilute screening power. Our platform nominated cytokine receptor subunits for three of the four cytokines present in our culture media as positive (IL-2, IL-7) or negative (IL-21) regulators of CAR T cell persistence – and potentially a fourth (IL-4) under supraphysiological levels of STAT6. These results suggest, unsurprisingly, that fine details such as media composition are likely to heavily influence screening outcomes. Our proof-of-concept experiments summarized here hint at broader utility in the context of more complex screening landscapes. In the future, we envision sizable impacts to gene discovery when this technology is applied to long-term T cell cultures, particularly in the setting of serial T cell challenge or simulated tumor microenvironments, coupled with multi-phenotypic or functional readouts.

## MATERIALS AND METHODS

### Plasmids and vectors

Our SAM.1 (PB-UniSAM) parent vector was a gift from Lesley Forrester (Addgene 99866), that itself was a polycistronic derivative of Feng Zhang’s dual vector system (Addgene 61422, Addgene, 73797). Successive daughter plasmids were generated via simple ligation or Gibson assembly using annealed complementary oligonucleotides (Integrated DNA Technologies) or premade dsDNA templates (e.g. Integrated DNA Technologies, GeneArt, PCR-amplification sourced) using routine molecular biology methods. Constructs containing second-generation synergistic activation mediator components (e.g. PP7) were cloned utilizing synthesized templates derived from previously published sequences.^78,79^

To generate lentiviral transfer vectors, we deposited CRISPRa, CAR, and reference components within a third-generation version of pHIV7.^80^ To minimize construct size for T cell delivery, our CRISPR-CAR CDSs were cloned into the PiggyBac donor Nanoplasmid™ that features an RNA-OUT^81^ counterselection system (Aldevron). These donor plasmids were further modified with transposon-flanking HS-A insulator sequences^82^ and a miniaturized insulator sequence inserted between coding cassettes.^83^ The origin of CDSs is indicated in **table S1.**

For intronization studies, candidate introns were positioned within the mScarlet-dCas9 CDS using Intronserter^84^ and scored for high splicing fidelity *in silico* using NetGen2.^85^ Ultimately 11 native human introns were chosen based on their: (1) small size (<250 bp); (2) status as documented housekeeping, cyclin, tumor suppressor, or T cell-enriched genes; (3) invariant splicing across every known mature RNA isoform; and (4) high splice donor and acceptor NetGene2 scores (≥ 0.94) when positioned within the mScarlet-dCas9 CDS. Intron sequences can be found in **table S2**.

All cloned constructs were sequence-verified via end-to-end Sanger sequencing (Genewiz) and/or long-read Nanopore sequencing (Plasmidsaurus). BaEVRless and Measles H/F plasmids were the kind gifts of Els Verhoeyen (CIRI, Université Côte d’Azur). All arrayed gRNA sequences used in our study are listed in **table S3**.

### Subcloning gRNA-LV pooled libraries

The human Calabrese CRISPR activation pooled library was a gift from David Root and John Doench (Addgene 92379, 92380). To subclone this pool we digested our protospacer-empty Libr.1*-null* transfer vector with BsmBI-v2 (New England Biolabs, NEB) overnight at 55 °C. To generate a pool of inserts for Gibson assembly, we performed a series of separate identical PCR reactions to minimize jackpotting. Reaction mix included forward [5’- GGCTTTATATATCTTGTGGAAAGGACGAAACACCG-3’] and reverse [5’- TCCTTTCGGCCCAGCATAGCTCTTAAAC-3’] primers, Calabrese template, in 2X NEBNext® Ultra™ II Q5® Master Mix (NEB) subjected to the following reaction: 98 °C 30 sec; five cycles of 98 °C 10 sec, 54 °C 30 sec, 65 °C 30 sec; 20 cycles of 98 °C 10 sec, 60 °C 30 sec, 65 °C 30 sec; 65 °C 2 min; hold at 4 °C. The next day, we ran our linearized plasmid backbone and 83 bp PCR products on a 1% agarose gel and recovered each using the Gel and PCR Clean-up Kit (Takara) per the manufacturer’s instructions. Insert and vector were combined at a 5:1 ratio and held at 50 °C for 1 hour in NEBuilder® HiFi DNA Assembly Master Mix (NEB). Gibson assembly products were purified using the MinElute® PCR Purification Kit (Qiagen). Purified product was delivered to NEB® 10-beta Electrocompetent *E. coli* using the Micropulser (BioRad) with pulse setting Ec1. Multiple EPs were pooled, with reaction number scaled to ensure transformation efficiencies yielding >200 clones per gRNA on average. Following a 1-hour recovery at 37 °C, 250 rpm, bacteria were directly inoculated into large scale culture vessels containing LB-Kan (30 μg/mL) and grown up overnight. The next day these cultures were pelleted and immediately processed for plasmid extraction using the Nucleospin® Xtra Maxi Plus EF (Takara).

To assess the quality of the new Jensen-Calabrese plasmid pool, gRNA protospacer sequences were amplified using NEB Q5 hot start HiFi master mix (M0515S) and Illumina-compatible primers (**table S4**). After clean-up with AMPure XP Beads (Beckman Coulter), the sequencing library was quantified by qubit and diluted to 4 nM. The library was sequenced using paired-end 150 bp reads on an Illumina HiSeq with 25% PhiX spike-in. Genome-wide libraries were sequenced to a targeted depth of 1000-fold coverage. All reads were analyzed by PoolQ v3.2.2 (https://github.com/broadinstitute/poolq). Sequencing indicated recoveries of 99.997% (AUC = 0.62) and 99.986% (AUC = 0.64) of subcloned Calabrese half-library A (AUC = 0.59, Addgene) and B (AUC = 0.61, Addgene) respectively (**Fig S10i**). Jensen-Calabrese plasmid protospacer frequencies for half-library A and B can be found in **table S5**.

### Lentiviral production

HEK293T/17SF cells between passage 3 and 20 were cultured in Gibco FreeStyle™ 293 medium at 1.5 × 10^6^ to 2 × 10^6^ cells/mL, shaking at 100 - 120 rpm, 37 °C, 5% CO_2_. Plasmids were combined in Opti-MEM™ (1:20 final volume), added at a ratio of 6.25 µg transfer plasmid, 6.25 µg Gag-Pol, 2.5 µg Rev, and 5 µg VSV-G per mL cell culture. For the envelope series, envelope plasmid substituted for VSV-G at an equivalent concentration, with the envelope plasmid mass doubled when two envelope plasmids were tested. Separately, PEIpro® transfection reagent was added to Opti-MEM™ (1:20 final volume) at 40 µL/mL cell culture. Plasmid and PEI mixtures were vortexed separately, then combined and incubated undisturbed at RT for 15 - 30 minutes before being added to cells. After 24 hours, sodium butyrate was added at 1 µL/mL culture. After an additional 48 hours, culture was centrifuged at 1000 × *g* for 15 minutes. The supernatant was filtered through a 0.45 µm PVDF Stericup filter, then centrifuged at 10,000 × *g* for 4 hours at 4° C with 30% brake. The resulting cell pellet was resuspended in OptiMEM™ at 1 - 1.5 µL/mL initial culture volume. A list of viral packaging components can be found in **table S6**.

For functional titration, Jurkat, Clone E6-1 (ATCC) and Raji (ATCC) cell lines between passage 3 and 20 were centrifuged and resuspended at concentration of 4 × 10^5^ cells/mL in RPMI-1640 supplemented with 10% fetal bovine serum (FBS) and 5% L-glutamine (cRPMI). Lentivirus aliquots were thawed at room temperature and mixed thoroughly. Diluted virus was added to the first column of a 96 well plate as a 1:25 dilution in cRPMI, then serially diluted 1:2 in cRPMI over 11 total wells. Protamine sulfate was added to the cell suspension at 8 µL/mL, and the mixture added to the virus dilution series at 1:2. Plates were incubated at 37 °C, 5% CO_2_. After 18 - 24 hours, an equal volume of cRPMI was added to each well and the plates were incubated for an additional 4 days. Plates were then centrifuged at 500 × *g* for 3 minutes and media removed. Plate wells were washed in PBS supplemented with 2% FBS and 250 mg/L sodium azide, then fixed in 0.5% formaldehyde. Fixed cells were stored at 4 °C awaiting data acquisition. Data was collected using a BD LRS Fortessa Flow Cytometer and analyzed using BD FlowJo software. Sample populations were gated on non-debris, single cells; transduced cells were compared to a non-transduced mock to determine positivity by the Overton method.

### Primary T cell isolation

For isolation of and cryopreservation of T cells, fresh leukopaks from StemExpress were processed within 24 hours of collection. Donors were limited to non-smokers between the ages of 18 - 35 and with a BMI <29.9. CD8 cells were isolated by incubating PMBCs with 3 - 5 mL CD8-specific StraightFrom® or CliniMACS® magnetic beads (Miltenyi) for 30 minutes at 4 °C with gentle agitation, then sorted on a Miltenyi MultiMACS™ Cell24 Separator and washed with autoMACS® Running Buffer. The CD8 enriched population was counted and cryopreserved in CryoStor CS5 (StemCell). The remaining cells CD8-depleted were labeled, sorted, and cryopreserved as before with 3 - 5 mL CD4-specific StraightFrom® or CliniMACS® magnetic beads (Miltenyi). The CD4- and CD8-depleted PBMCs were washed three times in running buffer to deplete platelets, then cryopreserved in CryoStor CS5.

For isolation and use of fresh, unfrozen T cells, leukocytes were recovered from spent leukocyte reduction system chambers (LRSCs), a byproduct of donor platelet collection (Bloodworks Northwest). CD8^+^ and CD4^+^ T cells were isolated via sequential positive selection using the RoboSep™ cell separation system (StemCell). Live CD4- and CD8-depleted PBMCs of the negative fraction were further purified by Lymphoprep™ (StemCell) and thoroughly washed. All recovered cell fractions were rested overnight in EP-TICLE culture media consisting of X Vivo-15 (Lonza) supplemented with 2% KnockOut serum replacement (Gibco), 4.6 ng/mL (∼83 IU/mL) IL- 2 (StemCell), 20 ng/mL IL-4 (Miltenyi), 10 ng/mL IL-7 (Miltenyi) and 20 ng/mL IL-21 (Miltenyi) at 37 °C, 5% CO_2_ prior to use.

### Classic lentiviral transduction

CD4^+^ and CD8^+^ T cells were combined 1:1 and stimulated with CTS™ Dynabeads™ CD3/CD28 (Invitrogen) at a 1:1 ratio overnight in X Vivo-15 (Lonza) supplemented with 2% KnockOut serum replacement (ThermoFisher), 4.6 ng/mL (∼83 IU/mL) IL-2 (StemCell). The next day, the culture volume was reduced and fresh media containing protamine sulfate (40 μg/mL final) (Fresenius Kabi) was added. Viruses were thawed, vortexed and added to each corresponding well, calibrated to a MOI = 1 (0.5 - 11% v/v). Cells were returned to the incubator, and fresh media was added six hours later, doubling the culture volume. Four days later, Mtx was added to a final concentration of 100 nM and maintained throughout culture. Seven days following activation, Dynabeads™ were removed with DynaMag columns (Invitrogen). On Day 10, cells were counted and assessed via flow cytometry. Cells were maintained in their selective media formulations throughout the duration of culture.

### Primary T cell EP and EP-TICLE

T cells were resuspended in supplemented P3 buffer (Lonza) with 1.5 μg (∼10 - 20 nM) PiggyBac donor plasmid and 40 nM RNA encoding PiggyBac transposase, generated via *in vitro* transcription (Aldevron). Cells were nucleofected using the Lonza 4D system with pulse program EO-115. For EP-TICLE experiments, electroporated cells were transferred to a G-REX culture vessel (Wilson Wolf) seeded with CD4- and CD8-depleted PBMCs and fresh EP-TICLE media. Mtx (Accord Healthcare) was added to cultures to a final concentration of 100 nM on Day 0 (fresh-sourced cells) or Day 3 (cryopreserved cells) and maintained throughout manufacturing. Media was replaced twice weekly, and cells were expanded for up to 21 days.

For conventional EP experiments, nucleofected primary T cells were transferred to G-REX wells with culture media alone. A list of culture reagents can be found within **table S7**.

### Flow cytometry

For relative antigen quantification, cells were collected and washed with 1X PBS. Cells were stained with antibodies or reactive dyes resuspended in 1X PBS for 30 minutes at room temperature. This was followed by two additional washes. For methods including a biotinylated primary antibody, the cells underwent a 20-minute subsequent stain at room temperature with Streptavidin-PE (Biolegend), followed by two additional washes. Following the last wash, the cells were fixed in 0.5% paraformaldehyde / 1X PBS. Cells were stored at 4 °C while awaiting data acquisition.

Data was collected using a BD LRS Fortessa Flow Cytometer. Data was analyzed using BD FlowJo software. Sample populations were routinely gated on SSC-A × FSC-A for non-debris and FSC-H × FSC-A for single cells. Cell positivity and intensity calculations were performed by FlowJo software. All antibodies and reagents used for flow cytometric analysis can be found in **table S8**.

### Duplex-droplet digital PCR (ddPCR)

Genomic (g)DNA of CRISPRa-CAR T cells from multiple donors was extracted at various timepoints throughout EP-TICLE manufacturing using the PureLink gDNA Mini Kit (Invitrogen). gDNA yield and purity were evaluated with the NanoDrop One spectrophotometer (Thermo Scientific). Prior to each experiment, fresh working stocks of gDNA were prepared by diluting with nuclease-free water to a concentration of 25 ng/µL.

For each 20 μL ddPCR reaction, 100 ng of gDNA was directly digested with 5 U of EcoRI-HF (New England Biolabs) in the ddPCR Supermix for Probes (no dUTPs) (Bio-Rad), with FAM/HEX primers and probes at final concentrations of 900 nM and 250 nM, respectively. Following droplet generation with the Automated Droplet Generator (Bio-Rad), PCR was performed in a C1000 Touch Thermal Cycler equipped with a 96-deep well reaction module (Bio-Rad). The PCR cycling program was optimized for the specific primer/probe sets and is as follows: 95 °C for 10 minutes (ramp rate of 2 °C/second), 40 denaturation/annealing cycles of 94 °C for 30 seconds (ramp rate of 2 °C/second) and 59.4 °C for 2 minutes (ramp rate of 1 °C/second), and a final extension at 98 °C for 10 minutes (ramp rate of 2 °C/second). Droplet plates were subsequently loaded into the QX200 Droplet Reader (Bio-Rad) for fluorescence analysis. Amplitude thresholds used to define positive and negative droplets were manually adjusted in the complementary QuantaSoft Analysis Pro software (Bio-Rad).

All primers and FAM/HEX-labeled probes were designed with PrimerQuest (Integrated DNA Technologies) and are listed in **table S9**. Primer/probe sets specific to the mScarlet, dCas9, or FMC63scFv CDSs were used to assess relative VCN of the CRISPRa transgene in relation to the CD19CAR transgene. Albumin (ALB) served as the reference gene for absolute VCN determination.

### Sequence analysis of plasmid donor plasmids

Long reads reported by Plasmidsaurus were aligned to each ddPCR probe string using minimap2.^86^ Using SAMtools^87^, a .sam file was converted to a .bam file in order to read out alignments. True proportions of each ddPCR probe string were calculated by taking the ratio of the total reads for each indicated sequencing string.

### Distal lentiviral transduction

Day 17 EP-TICLE CAR T cells were counted, spun down, and resuspended in complete TICLE media with 40 μg/mL protamine sulfate at a density of approximately 5 × 10^6^/mL. For gRNA-LV tiling, 10^7^ cells were transduced using 2.5% v/v concentrated gRNA-LV, resuspended in a final volume of 2 mL; spinoculation consisted of a 30 min centrifugation at 800 × *g*, 32 °C, low brake. Immediately thereafter, cells were returned to the incubator. Approximately 6 hours later, cultures were volumed up to 4 mL using fresh TICLE media. For medium-scale culture experiments, cells and viruses were scaled proportionally.

For genome-wide screening, Day 16 EP-TICLE CAR T cells (**Fig S13b**) were transduced at 0.625% v/v (MOI ≈ 0.3 - 0.6), plated at 5.2 × 10^6^/cm^2^; donor-specific gRNA library functional titers were pre-determined via a pilot study with identical donor apheresis cells (**Fig S13a**). Two days after transduction, cells were selected with puromycin at 2.0 μg/mL for 48 hrs and then assessed for gRNA positivity as indicated by copGFP. Due to lower-than-expected purification efficacy (**Fig S13c**), selection was extended for an additional day at reduced cell density (<4 × 10^6^/cm^2^). Cell positivity was reassessed on landscape Day 3 and observed to have markedly increased (**Fig S13d**).

### Genome-wide CRISPRa-CAR T cell persistence screen

Following cell manufacturing and selection, a culture cell sample was taken (Day 0), washed, and frozen down for later processing. The next day, CRISPRa-CAR-library^+^ cells were equally distributed across three treatment arms: (1) a 1:1 (E:T) Raji co-culture in TICLE media, (2) a 1:1 (E:T) Raji co-culture in cytokine-free media, or (3) TICLE media alone. Cells were fed and counted every 3 - 4 days, at which point, complete media exchanges also occurred. Culture samples were banked on Day 7, as cells in the cytokine-negative conditions dwindled (**Fig S13e**). At all timepoints and across all treatment arms, culture samples were maintained at sufficient numbers (≥ 26 × 10^6^) to maintain coverage of > 400 copies per gRNA on average. Cells in the cytokine-free arm had a high incidence of death, as expected, and were processed via the Dead Cell Removal Kit (Miltenyi) prior to freezing. Upon observing a higher-than-anticipated proportion of viable cell loss following dead cell removal, only Donor A met predetermined cell count cutoffs for subsequent sequencing.

### Protospacer gDNA amplification and sequencing

To process gDNA, banked cell pellets were rapidly thawed and processed using the Nucleospin® Blood XL kit (Takara) following the manufacturer’s instructions. gDNA quantity and quality was assessed using the NanoDrop® One (Thermo Scientific). For protospacer amplification, gDNA was diluted 1:4 with molecular grade H2O to reduce the effect of PCR inhibitors. gDNA template was combined with Illumina-compatible (0.5 μM) forward and (0.5 μM) reverse primers, dNTPs, 10X buffer, and Ex Taq polymerase (Takara) per the manufacturer’s instructions. Combined gDNA template and PCR master mix was divided into multiple 100 µL reactions to reduce the effect of PCR jackpotting. Thermocycling took place in a deep well plate (BioRad), with conditions consisting of an initial denaturation of 95 °C; then 28 cycles of 95 °C 30 sec, 53 °C 30 sec, 72 °C 30 sec; a final extension of 72 °C for 10 min; and holding at 4 °C. Following amplification, each 100 µL reaction was re-pooled. Each sample was run on a 4% agarose gel and TapeStation 4150 (Agilent) to verify the presence of PCR products of the appropriate size (383 bp). A fraction of each sample’s pooled PCR product was cleaned using AMPure XP beads (Beckman Coulter), then quantified by qubit and diluted to 4nM. Multiple samples were pooled, and each sequencing library was Illumina-sequenced as described further above.

### CRISPRa-CAR screen analysis

For analysis of recovered libraries, reads were aligned to the combined Calabrese A and B pools as a single reference file using MAGeCK version 0.5.9.4.^40,88^ Each of the matching samples across library A and B were merged to generate a single normalized read count table. Normalized read counts in high versus low bins were compared using mageck test with –norm-method none, –paired options, pairing samples by donor. Genes with a false discovery rate (FDR) <0.05 were identified as positive (enriched) or negative (depleted) “hits.” gRNA frequencies from all screening arms can be found in **tables S10 - S12**. These normalized read count tables were entered into R version 4.2.1. for data visualization. Gene set enrichment analysis (GSEA) was performed using the same normalized read count table, an output of mageck. Genes classified as negative hits in **Fig 4b** were imported into a WEB-based GEne SeT AnaLysis Toolkit (WebGestalt)^89^ The gene list was filtered for annotated regulators of apoptosis and ranked from most- to least-depleted using log10(negative.score) (**Fig 4f**). To calculate general screen sensitivity, non-redundant GOBP pathways with genes having the most negative normalized enrichment scores (NESs) were ranked and plotted (**Fig 4g**). A full ranking of these genes can be found in **table S13**.

Correlation coefficients among sequenced samples were calculated through R cor() function (**Fig. S14c**). To calculate significantly enriched pathways, EnrichAnalyzer from the mageckflute package was used to identify enriched molecular pathways across negatively and positively enriched genes from each screen. Reference pathways were drawn from KEGG, REACTOME, and C2_CP databases; and were annotated if within at least one screen, they satisfied all of the following: NES > abs(0.5), FDR < 0.05, p.adj < 0.01 (**Fig S14f**). A full list of significant pathways, broken out by treatment arm, is included in **table S14**.

### Cell killing assay

For T cell cytotoxicity assays, CRISPRa-CAR T cells generated from EP-TICLE and transduced with gRNA-LV were plated 1:1 with mCherry^+^ Raji targets in Nunc™ Edge™ flat-bottom microplates. Whole wells were imaged using the Incucyte® Live-Cell Analysis System (Sartorius) every four hours up until 30 hours had passed. Total target cell counts were obtained for each timepoint using the Incucyte® Basic Analyzer tool and normalized to time zero in the absence of T cell effectors.

### Immortalized cell culture EP and maintenance

Jurkat and Raji cell lines were cultured in RPMI-1640 (Gibco) supplemented with 10% FBS (Hyclone) and 1% 200 mM L-glutamate (Gibco). Cells were passaged 1 or 2 times a week, split 1:5 or 1:10, respectively. Cells were not allowed to grow denser than 1.5 × 10^6^/mL. For electroporation, cells were split the day before manipulation to ensure high viability and maximize EP efficiency. For transient transfection experiments, 1 - 2.5 × 10^6^ Jurkat T cells were resuspended in EP buffer from the Nucleofector® Kit V (Lonza). Cells were combined with 1.5 μg of donor plasmid in a reaction volume of 100 μL and electroporated on the Amaxa® Nucleofector® II using pulse code X-005.

## Supporting information

Supplemental Results

## Supplemental Figures

**Figure S1:**
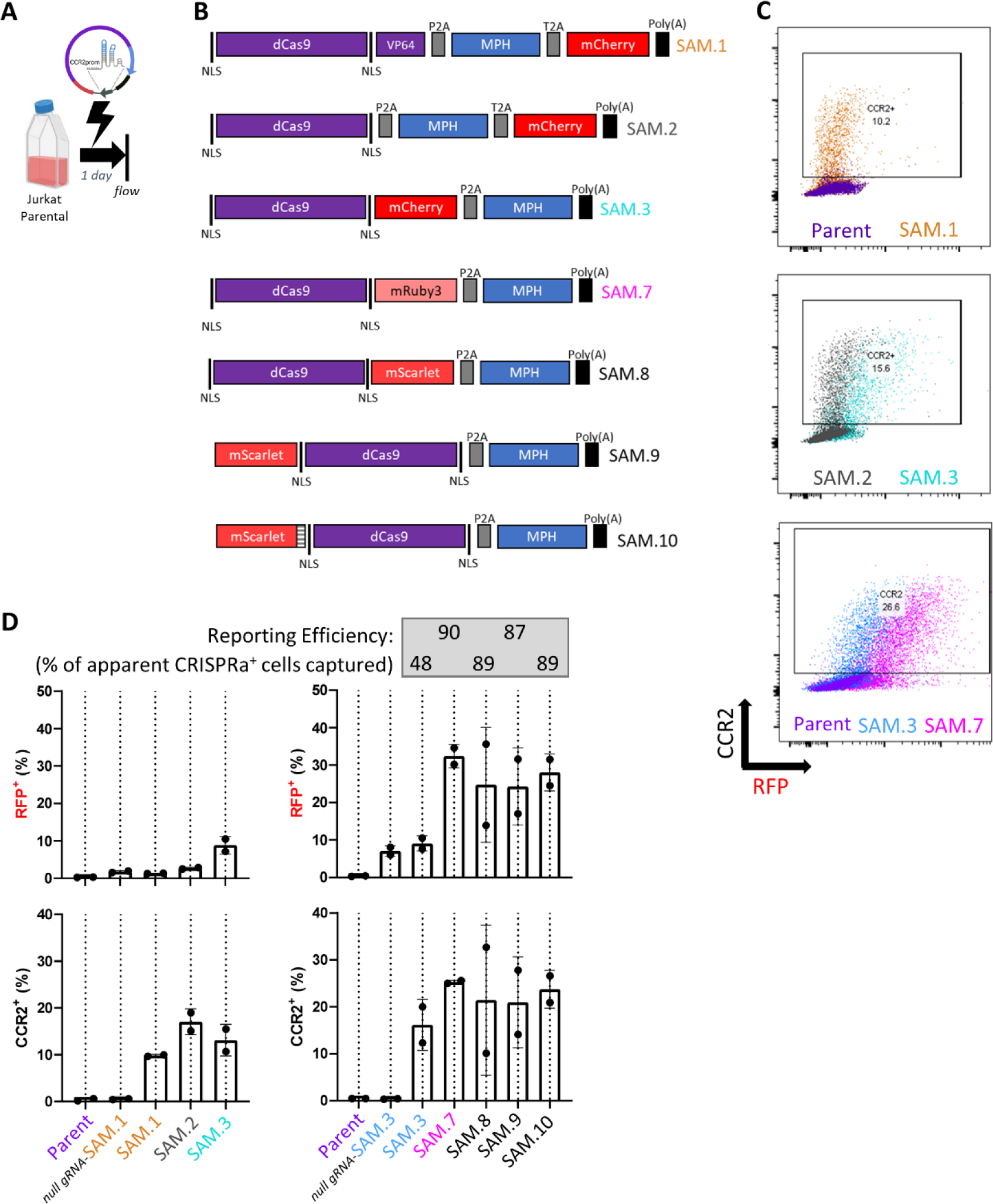
Optimization of CRISPRa Reporter. (**A**) General schema of transient delivery of CRISPRa CCR2 overexpression plasmids to Jurkat T cells. (**B**) CRISPRa fluorescent reporter variant designs. (**C**) Select flow cytometry plots and (**D**) quantification of CCR2 (CRISPRa target gene) and red fluorescent reporter among indicated designs from (A) delivered transiently to Jurkat T cells via electroporation with plasmid and assessed the next day. All quantified data was obtained via flow cytometry gated on live singlets. Reporting efficiency indicated in (D) is calculated as %CCR2^+^ cells / %RFP^+^ cells. Quantitative data in (D) depicts mean and standard deviation of three EP replicates.

**Figure S2:**
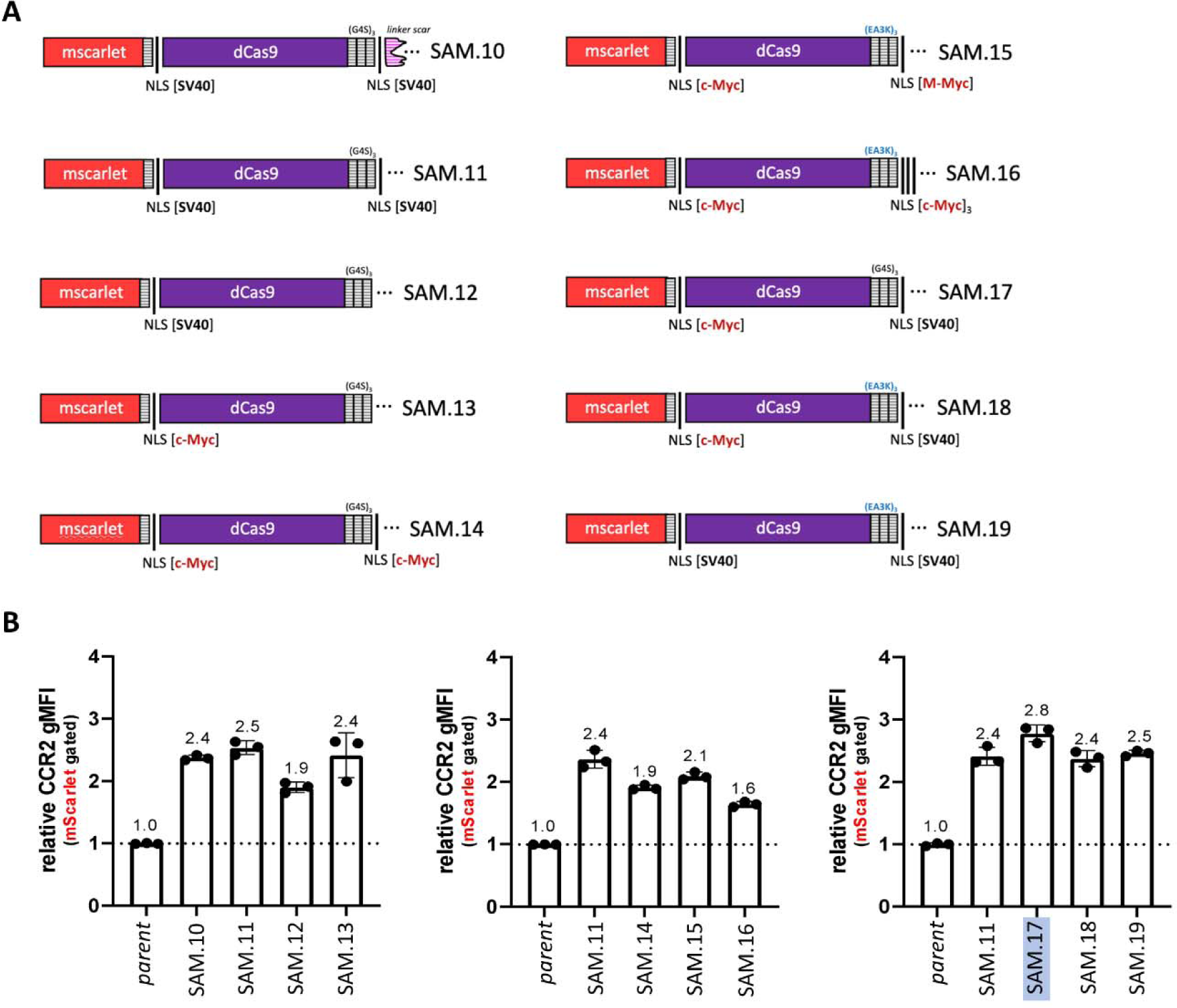
Optimization of dCas9 NLS. (A) NLS variant designs. (B) Quantification of CRISPRa endogenous target (CCR2) positivity across NLS designs transiently delivered via electroporation using plasmid Jurkat T cells and assessed the next day. All quantified data was obtained via flow cytometry gated on mScarlet-dCas^+^ live singlets or live singlets only (parent cultures). Quantitative data in (B) depicts mean and standard deviation of three EP replicates.

**Figure S3:**
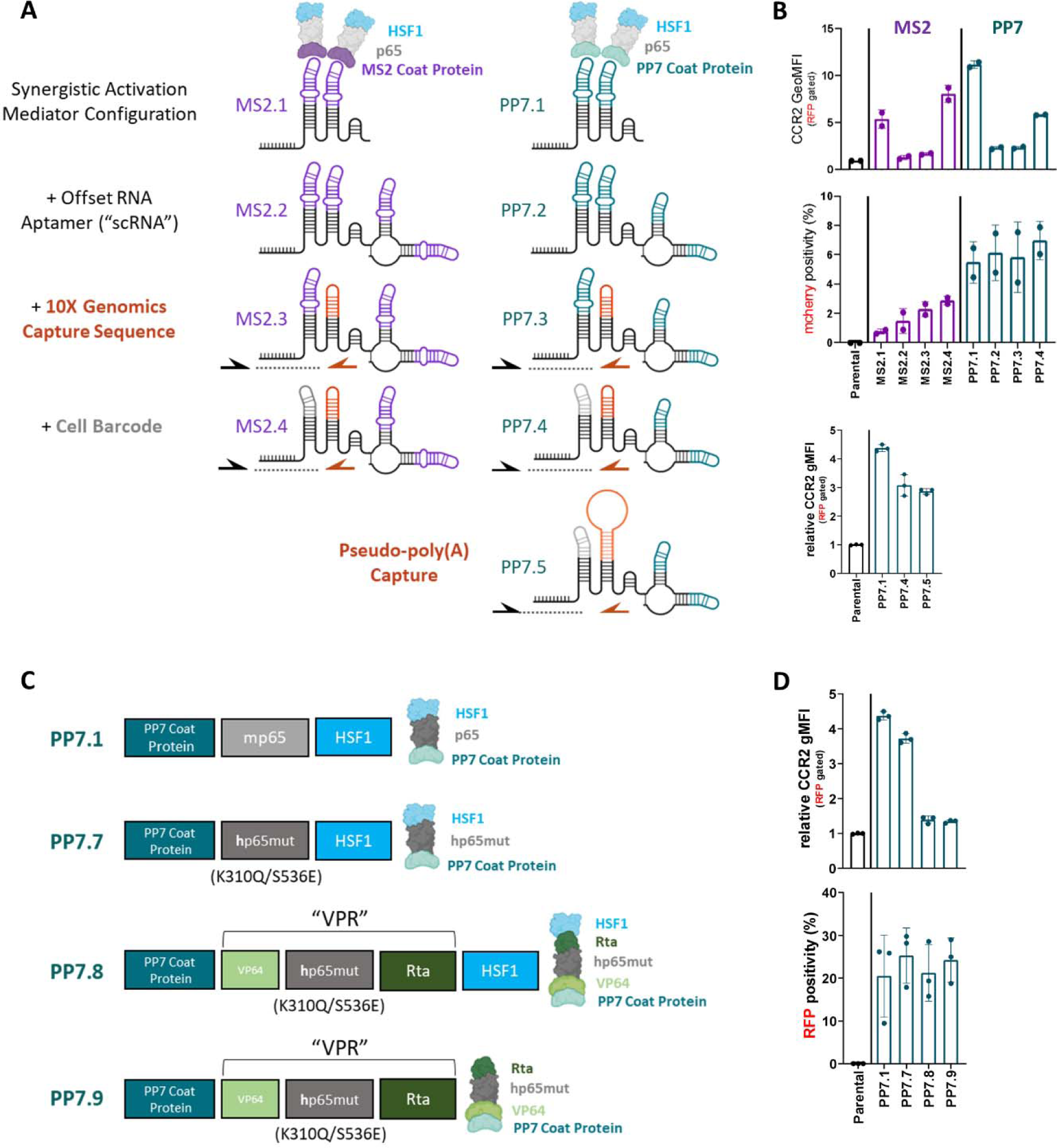
Optimization of CRISPRa aptamer-activator and gRNA variants. (A) gRNA variant designs. (B) Quantification of CRISPRa endogenous target (CCR2) (*top and bottom graphs*) and fluorescent reporter (mcherry) (*middle graph*) positivity across gRNA designs transiently delivered via electroporation using plasmid Jurkat T cells and assessed the next day. (C) PP7 coat protein CRISPRa effector variants designs. (D) Quantification of CRISPRa endogenous target (CCR2) (*top graph*) and fluorescent reporter (*bottom graph*) positivity across gRNA designs transiently delivered via plasmid to Jurkat T cells via electroporation and assessed the next day. All quantified data was obtained via flow cytometry gated on live singlets. Quantitative data in (B) and (D) depict mean and standard deviation of two or three EP replicates.

**Figure S4:**
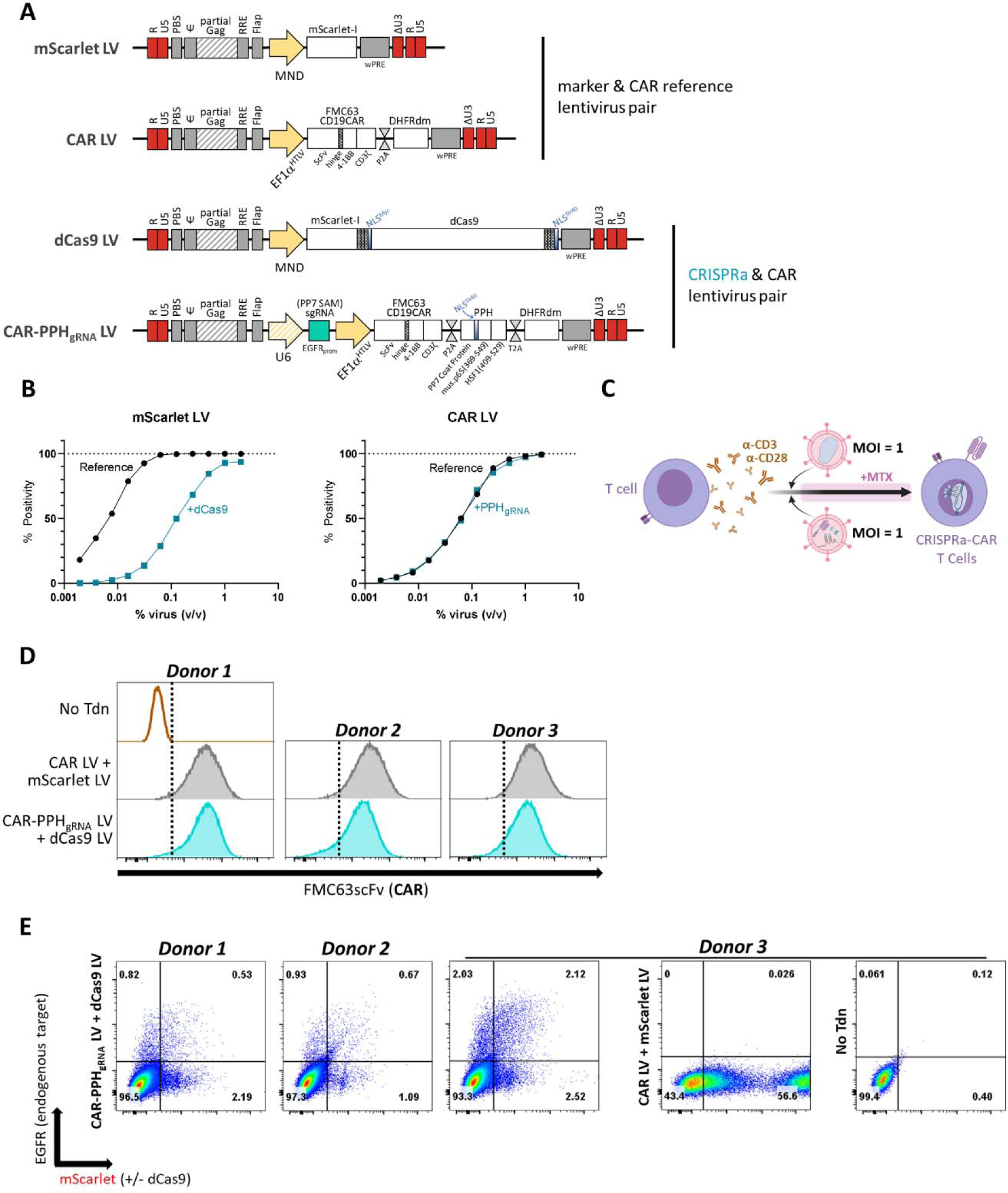
CRISPRa-CAR T cell production via lentiviral transduction. (A) Lentiviral vRNA payload maps of CRISPRa/CAR pair and reference pair. (B) Plots depicting titration of CRISPRa lentiviruses and reference lentiviruses in Jurkat T cell line. (C) Cell production schema of α-CD3/α-CD28 bead-activated (Day 0) CD8 and CD4 T cells transduced (Day 1) with CRISPRa/CAR or reference lentivirus pairs at an MOI = 1 per each virus, then MTX selection initiated (Day 4) and maintained throughout. (D) Lentiviral cell manufacturing Day 7 flow cytometry histograms of FMC63scFv (CAR) expression from T cell populations generated via mScarlet-LV/CAR-LV or dCas9-LV/CAR-PPH_gRNA_ lentivirus pairs. (E) Flow cytometry plots of mScarlet(+/−dCas9) and EGFR expression from cells described in (D). Cells in (D) and (E) are gated on CD8^+^ or CD4^+^ live singlets.

**Figure S5:**
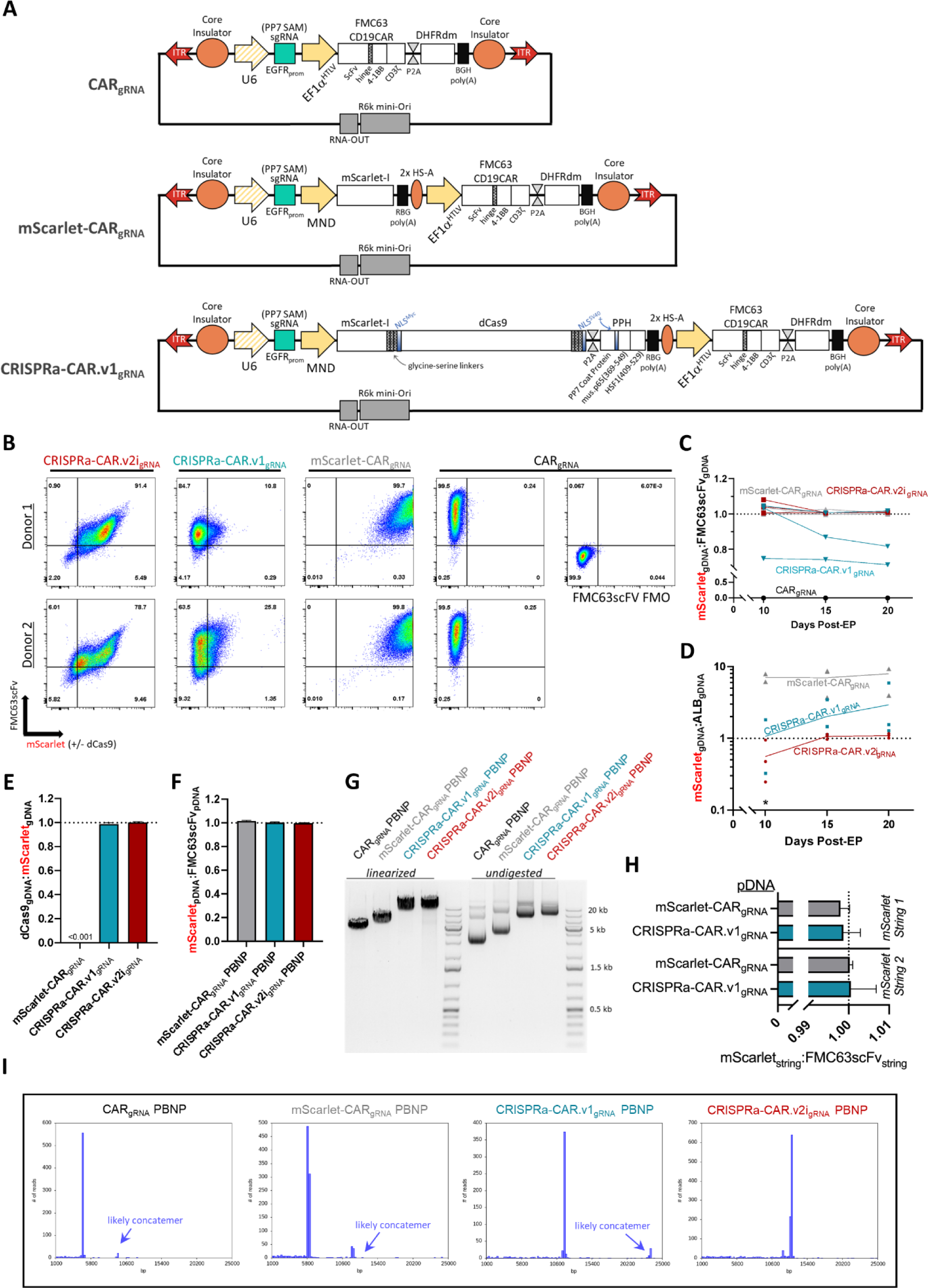
Extended flow cytometry of CRISPRa-CAR T cells, donor plasmid maps and analysis. (A) PiggyBac nanoplasmid donor maps. (B) Extended flow cytometry plots of FMC63scFv (CAR) and mScarlet (+/−dCas9) expression in CRISPRa-CAR.v2i_gRNA_, CRISPRa-CAR.v1_gRNA_, mScarlet-CAR_gRNA_, and CAR_gRNA_ mixed CD8^+^ and CD4^+^ cells at Day 20 of EP-TICLE. (C) Droplet-digital (dd)PCR relative quantification of mScarlet_gDNA_ to FMC63scFv_gDNA_ during EP-TICLE manufacturing in cells generated using CAR_gRNA_, mScarlet-CAR_gRNA_, CRISPRa-CAR.v1_gRNA_, or CRISPRa-CAR.v2i_gRNA_ donor plasmid. (D) ddPCR absolute quantification of mScarlet_gDNA_ to Albumin (ALB)_gDNA_ during EP-TICLE manufacturing in cells generated using mScarlet-CAR_gRNA_, CRISPRa-CAR.v1_gRNA_, or CRISPRa-CAR.v2i_gRNA_ donor plasmid. (E) ddPCR relative quantification of dCas9_gDNA_ to mScarlet_gDNA_ as a function of indicated donor plasmid. (F) ddPCR relative quantification of mScarlet_gDNA_ to FMC63scFv_gDNA_ detection in mScarlet-CAR_gRNA_, CRISPRa-CAR.v1_gRNA_, and CRISPRa-CAR.v2i_gRNA_ donor plasmid pools. (G) Visual agarose gel electrophoresis of CAR_gRNA_ (4.8 kb), mScarlet-CAR_gRNA_ (6.0 kb), CRISPRa-CAR.v1_gRNA_ (11.8 kb), and CRISPRa-CAR.v2i_gRNA_ (13.0 kb) donor plasmids with (*left*) or without (*right*) linearization via NruI restriction enzyme. (H) Nanopore sequencing DNA string read frequency of mScarlet_gDNA_ and FMC63scFv_gDNA_ ddPCR amplicons for CRISPRa-CAR.v1_gRNA_ donor plasmid. (I) Representative nanopore sequencing read length reports for each donor plasmid with putative multimers labeled. Each curve in (C) and is generated from cells sourced from a distinct blood donor. Each curve in (D) is the average of 2 - 3 individual donors (denoted by individual points). *samples did not have sufficient gDNA for analysis at Day 10 in (D).

**Figure S6:**
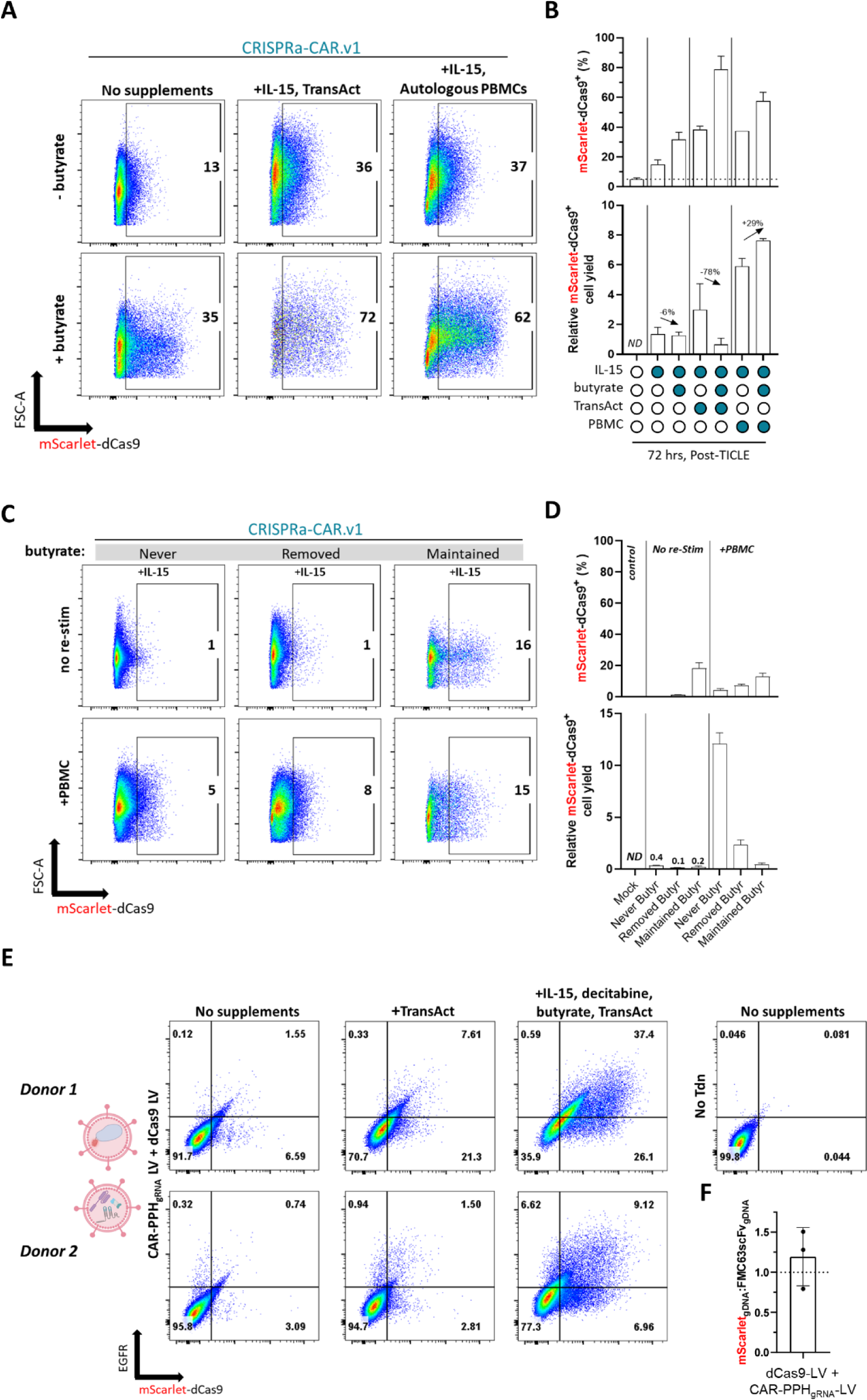
Extended de-silencing of CRISPRa-CAR T cells generated via transposon-based or lentiviral integration methods. (A) Flow cytometry plots and (B) accompanying quantification of mScarlet-dCas9 cell positivity in CRISPRa-CAR.v1 CD8^+^ T cells generated via EP-TICLE and treated with various de-silencing compounds for 72 upon completion of cell manufacturing. (C) Flow cytometry plots and (D) accompanying quantification of mScarlet-dCas9 cell positivity in CRISPRa-CAR.v1 CD8^+^ T cells generated via EP-TICLE and treated thereafter with or without autologous PBMCs and variable butyrate exposure to include 10 days off (“never”), 4 days on/ 6 days off (“removed”), or continuous exposure for 10 days (“maintained”). (E) Flow cytometry plots of mScarlet (+/−dCas9) and EGFR expression in mixed CD4^+^ and CD8^+^ T cells transduced with CRISPRa and CAR lentiviral pairs (vector maps shown in Fig 3.1a) and treated with various de-silencing compounds for 48 hours upon completion of cell manufacturing after 11 days. (F) ddPCR relative quantification of mScarlet_gDNA_ to FMC63scFv_gDNA_ in cells generated via transduction with dCas9-LV/CAR-PPH_gRNA_ lentivirus pairs. Cells for (A thru D) were sourced from a single blood donor but (B and D) represent independent EP-TICLE cultures (n = 2) per condition shown. Cells in (A) thru (E) are gated on live singlets.

**Figure S7:**
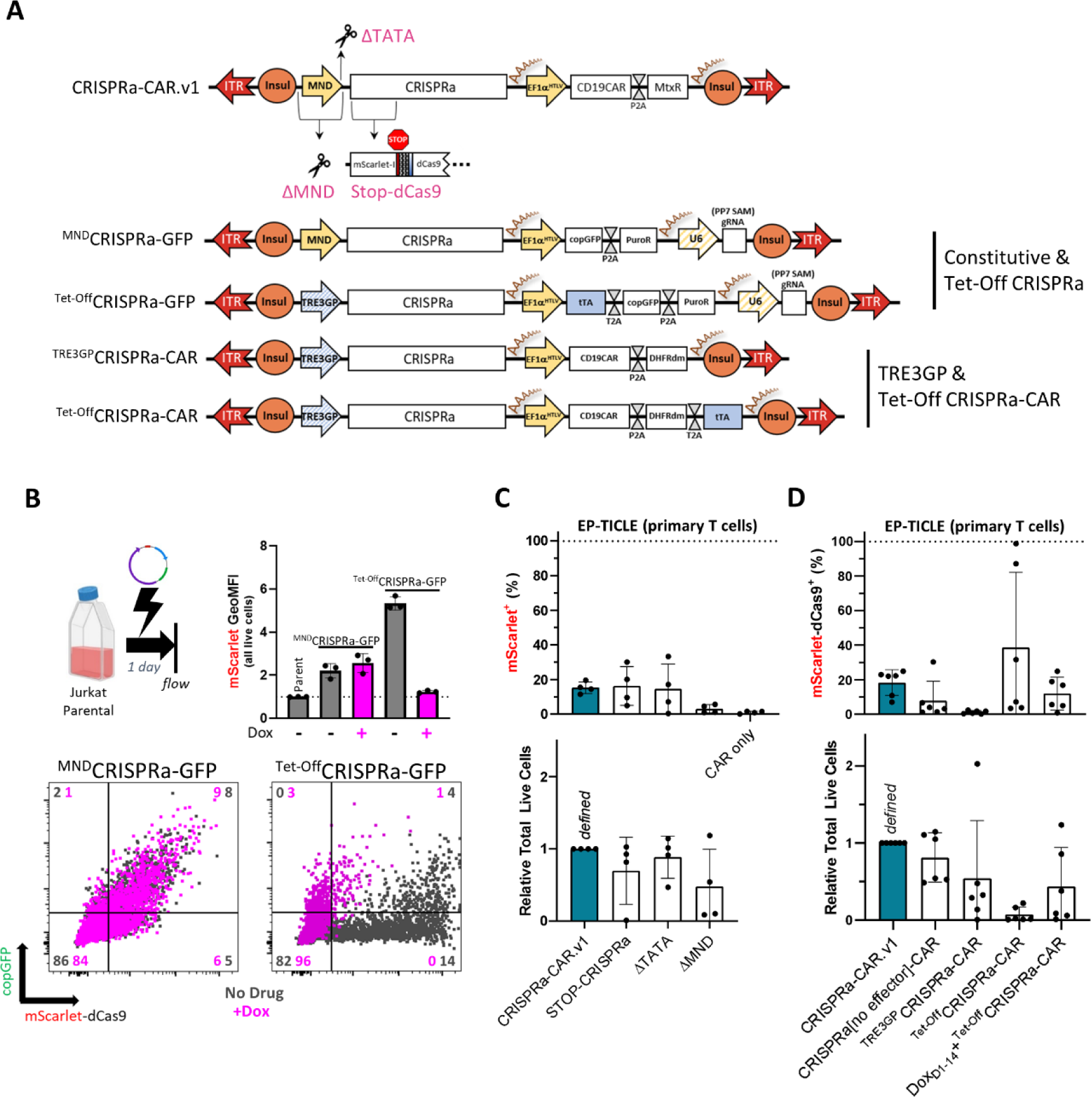
Restrained and modified CRISPRa expression attempts. (A) CRISPRa and CRISPRa-CAR plasmid donor maps. (B) Experimental setup (*upper left*), mScarlet-dCas9 geometric mean fluorescence intensity (*upper right*) and flow cytometry plots of copGFP and mScarlet-dCas9 expression (*bottom*) in Jurkat T cells transiently electroporated with donor plasmids depicted in (A) with or without doxycycline (dox) treatment. (C and D) Quantification of mScarlet(+/−dCas9) positivity (*top graph*) and relative total live cells (*bottom graph*) of 1:1 CD4 and CD8 bulk EP-TICLE cultures made from indicated donor plasmids. All cells are gated on live singlets. Calculated values in (C) and (D) are generated from data additionally gated on CD4^+^ or CD8^+^ cells, generated from 4 or 3 (where cells limiting) human blood donors.

**Figure S8:**
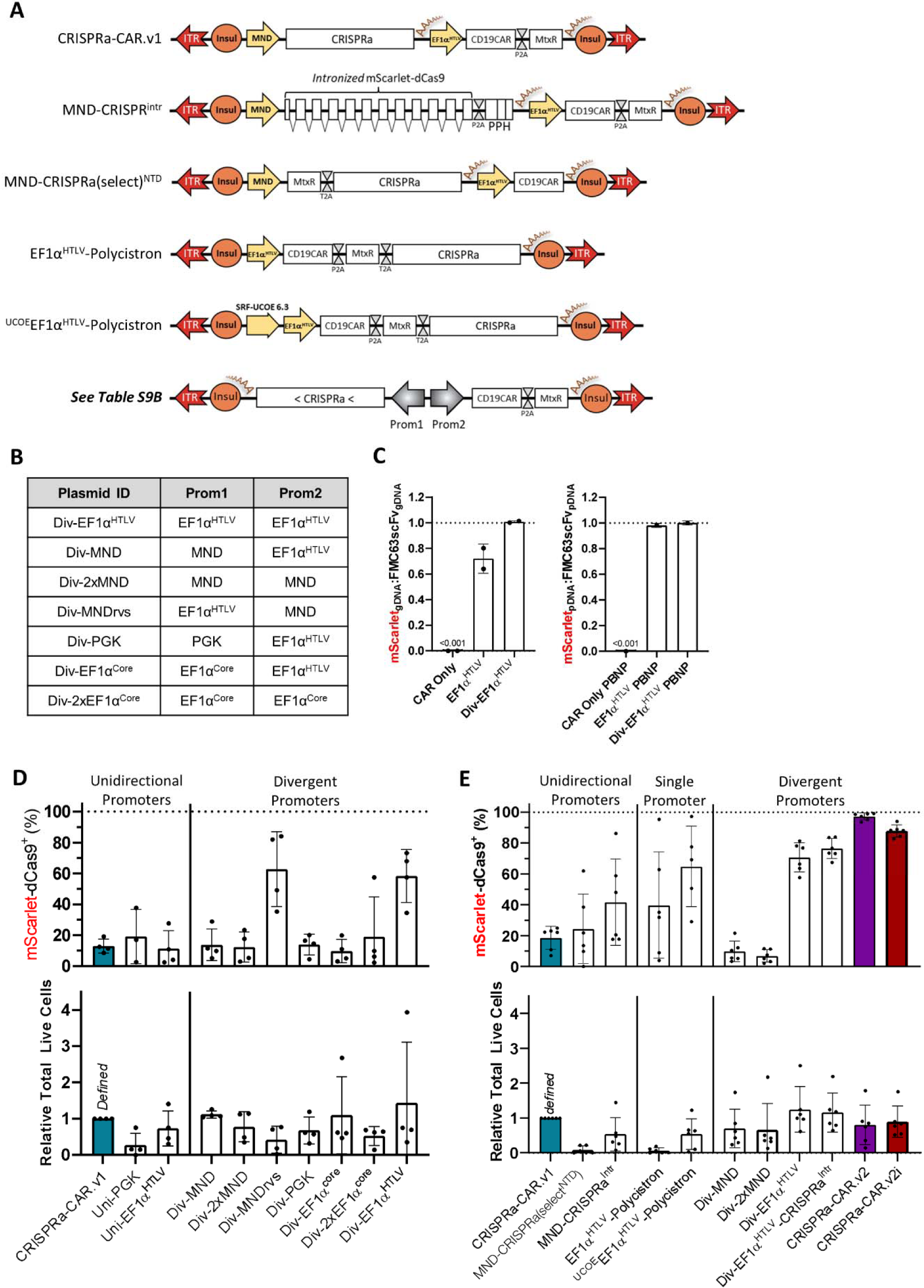
Partial or variant CRISPRa-CAR vector optimization. (A) Abridged donor plasmid maps of select CRISPRa-CAR variants and (B) Accompanying table outlining indicated divergent promoter pairs. (C) Ratio of T cell mScarlet_gDNA_ / CAR_gDNA_ as a function of time and EP-TICLE donor plasmid indicated in plot (*left*) and ratio of mScarlet_pDNA_ / CAR_pDNA_ measured in donor plasmid prep (*right*). (D) Ratio of T cell mScarlet_gDNA_ / CAR_gDNA_ as a function of time and composition of EP-TICLE donor plasmid admixtures indicated. (E and F) mScarlet-dCas9 positivity (*top graph*) and relative total live cell yield (*bottom graph*) as a function of EP-TICLE donor plasmid. mScarlet-dCas9 positivity is measured by flow cytometry, gated on CD4^+^ or CD8^+^ live singlets. Cells in (D) and (E) are gated on CD8^+^ or CD4^+^ live singlets only. All experiments depicted here used 1:1 mixtures of CD8^+^ or CD4^+^ T cells as EP-TICLE input. Quantified data in (D and E) and compiled from cells generated from 3 (where cells limiting) to 6 human donors. Data depicted in (D) and (E) originated from independent experiments.

**Figure S9:**
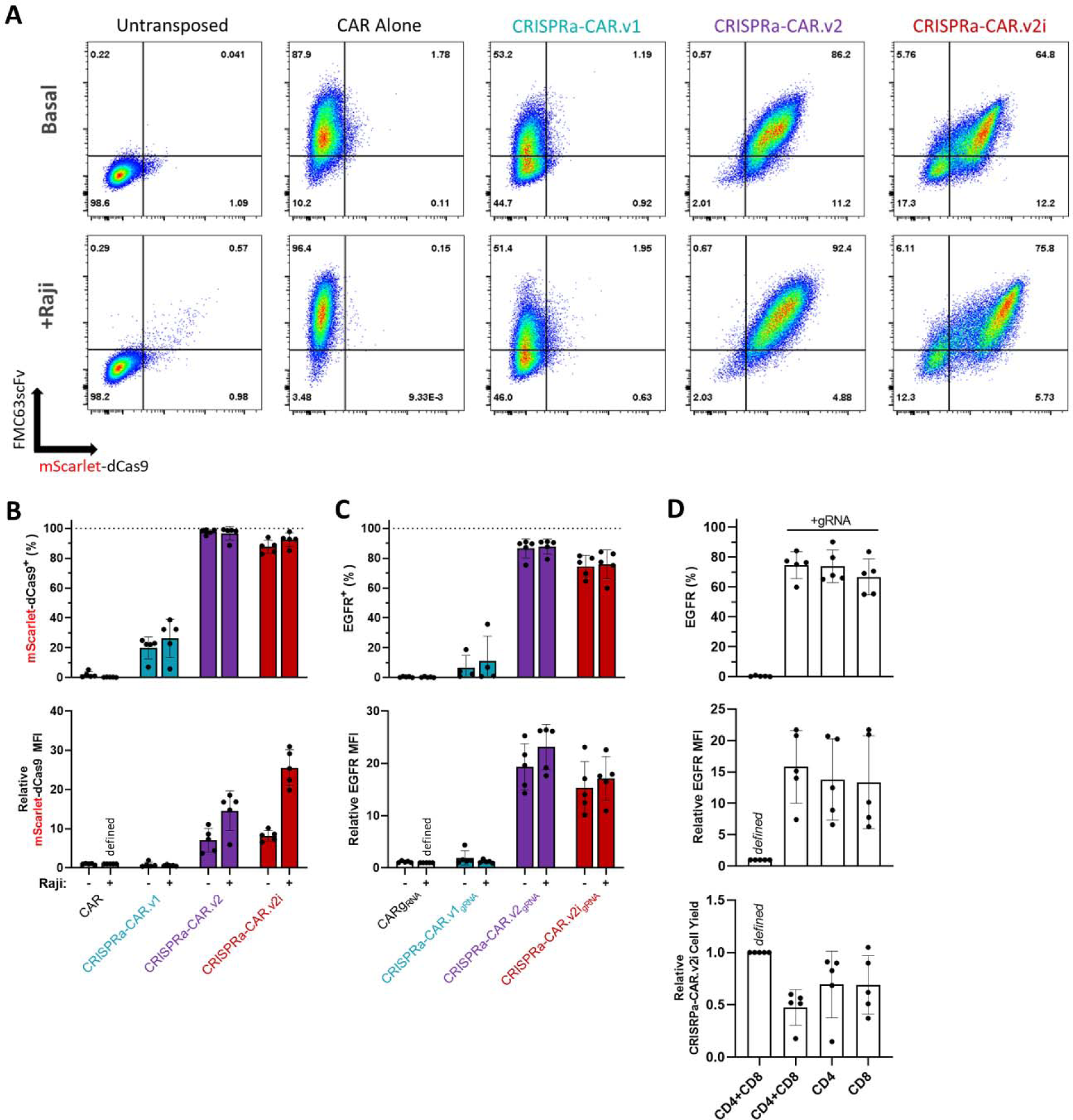
Additional analyses of top candidate CRISPRa-CAR T cells produced via EP-TICLE. (A) Day 20 flow cytometry plots of FMC63scFv mScarlet-dCas9 expression in cells generated via EP-TICLE from indicated donor plasmid (or mock transposition, TransAct-stimulated control cells), with or without 48-hr Raji cell line culture at a 1:2 effector-to-target ratio. (B) Quantification of Day 20/21 mScarlet-dCas9 positivity (*top graph*) and relative mScarlet-dCas9 median fluorescence intensity (*bottom graph*) of indicated CAR T cells generated via EP-TICLE. (C) Quantification of Day 20/21 EGFR positivity (*top graph*) and relative EGFR median fluorescence intensity (*bottom graph*) of indicated CAR T cells generated via EP-TICLE. (D) Quantification of Day 20/21 EGFR positivity (*top graph*), relative EGFR median fluorescence intensity (*middle graph*), and relative mScarlet-dCas9^+^ cell yield *(bottom graph)* of CRISPRa-CAR.v2i T cells generated from indicated starting cell T cell population via EP-TICLE. Unless otherwise labeled, all experiments depicted here used 1:1 mixtures of CD8^+^ or CD4^+^ T cells as EP-TICLE input. Cells in (A) thru (D) are gated on CD8^+^ or CD4^+^ live singlets.

**Figure S10:**
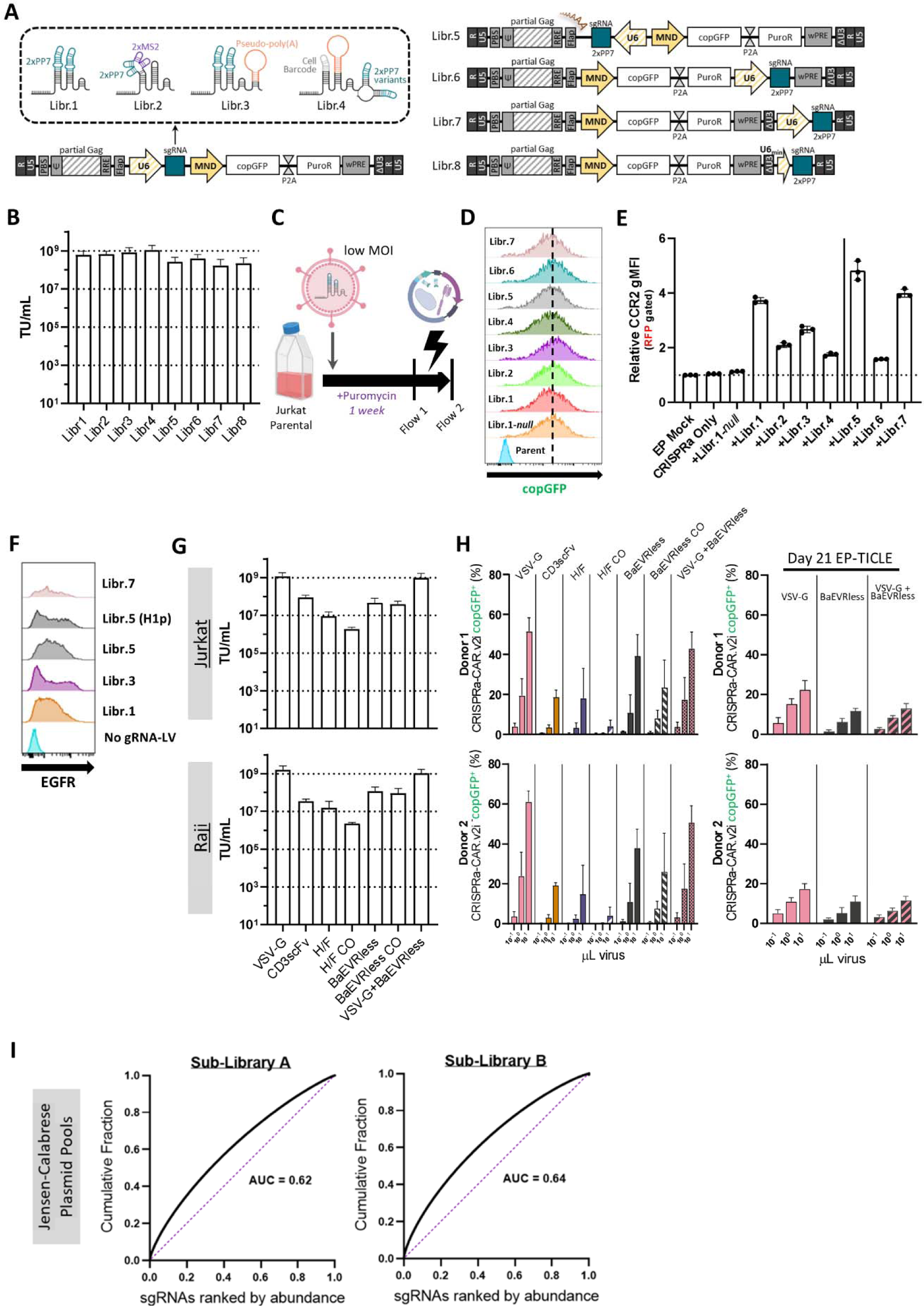
Additional analyses of gRNA-LV ssRNA genome and packaging envelope. (A) Abridged maps of library (libr.) series candidate vRNA lentiviral genomes. (B) Calculated lentiviral functional titers in Jurkat T cells of library LV series depicted in (A). (C) Graphic depicting experimental schema to generate and assess stable Jurkat cell lines. (D) histograms of copGFP expression assessed via flow cytometry of Jurkat T cell lines after puromycin selection at “Flow 1” timepoint. (E) Relative CCR2 geometric mean fluorescence intensity of indicated Jurkat parental (“EP mock,” “CRISPRa Only”) or puromycin-selected cell line the day after electroporation using CRISPRa-CAR.v1 PiggyBac donor plasmid (except “EP mock”). (F) Histograms of EGFR expression in CD8^+^ copGFP^+^ (except untransduced reference) lines as function of LV-gRNA species indicated. (G) Calculated lentiviral functional titers in Jurkat (*top graph*) and Raji (*bottom graph*) cell lines as a function of pseudotyped viral envelope (depicted in Fig 3A). (H) Extended figure depicting copGFP positivity of CRISPRa-CAR.v2i T cells as a function of viral packaging by designated viral envelope variant and volume of concentrated lentivirus, transduced on Day 17 (*left*) or 21 Day 21 (*right*) of EP-TICLE. (I) Jensen-Calabrese plasmid pool half-library A (*left*) and B (*right*) cumulative gRNA frequency as ranked by gRNA abundance. Area under the curve approximates library distortion (0.5 = perfect representation, 1.0 = perfect skew). Dashed purple lines in (I) indicate hypothetical AUC = 0.5. Cells in (F) are gated on mScarlet-dCas9^+^ live singlets. Cells in (G) were transduced on Day 17 of EP-TICLE and assessed for copGFP positivity on Day 21 after gating on live singlets.

**Figure S11:**
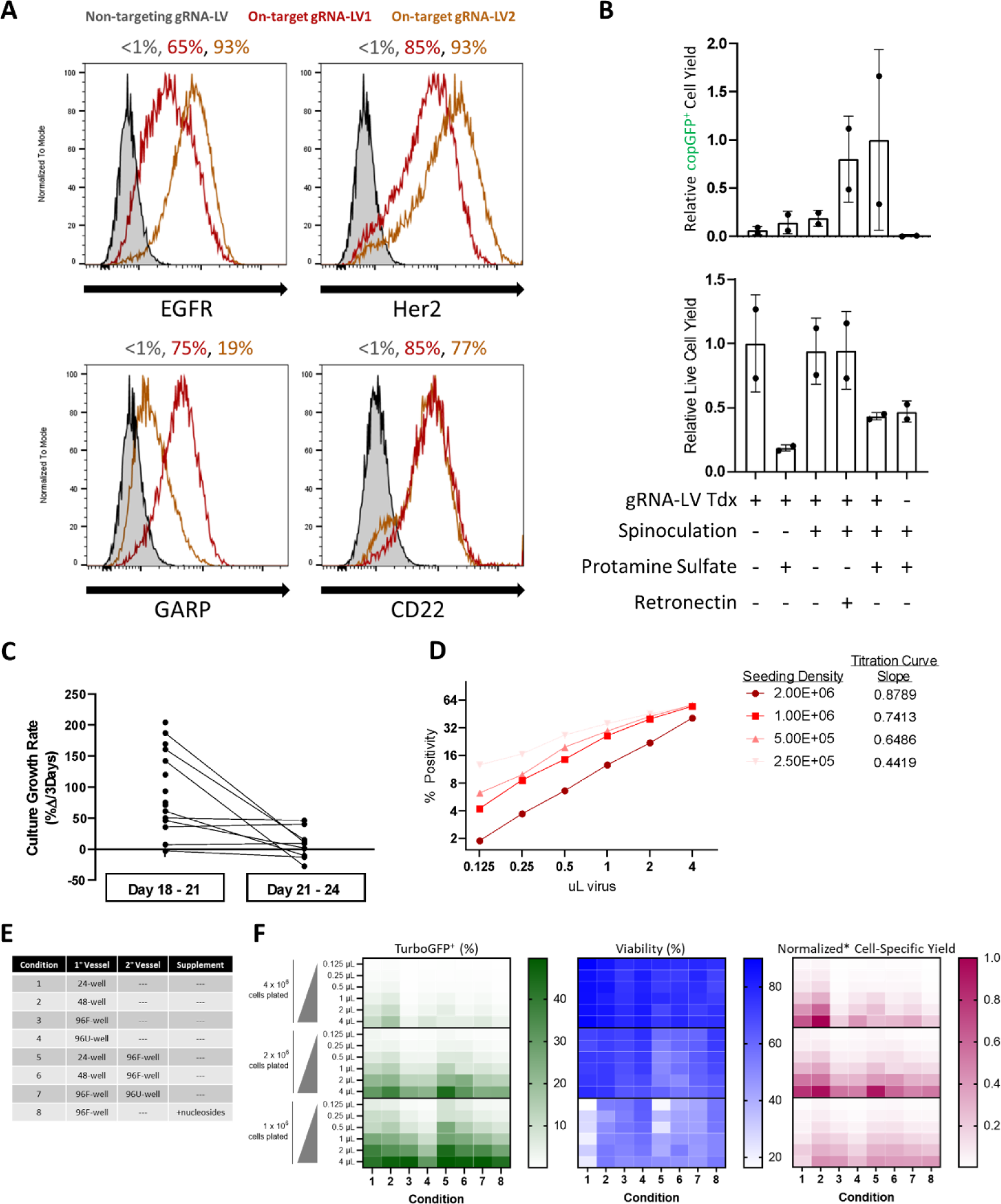
gRNA target expression and optimization of distal transduction conditions. (A) Representative histograms of indicated protein expression as determined by flow cytometry as a function of transduction of indicated gRNA-LV to CRISPRa-CAR.v2i T cells generated from single donor. (B) Calculated copGFP^+^ relative cell yield (*top graph*) and relative total live cell yield (*bottom graph*) relative to each indicated transduction condition in CRISPRa-CAR.v2i T cell generated by EP-TICLE. (C) Calculated growth culture growth rates of CRISPRa-CAR T cell candidate lines between EP-TICLE Day 18 thru 21 (*left*) and Day 21 thru 24 (*right*) spread across two human donors and CD8 only input cells. (D) Calculated copGFP positivity of transduced CRISPRa-CAR.v2i T cell transduced at Day 17 of EP-TICLE at four different cell densities and assessed four days later. (E) Table of transduction conditions profiled in figure (F). (F) Transduction copGFP positivity (*left heatmap*), cell viability (*middle heatmap*) and normalized cell-specific yield (*right heatmap*) of CRISPRa-CAR.v2i cells as a function of cell density, virus volume, plating conditions (*see bottom labels*). Cells in (A), (D), and (F) used 1:1 mixtures of CD8^+^ or CD4^+^ T cells as EP-TICLE input. Cells in (B) and (C) used CD8^+^ T cells as EP-TICLE input. All transductions were performed at Day 17 of EP-TICLE manufacturing and assessed four days later via flow cytometry. Cells in (A) are gated on mScarlet-dCas9^+^CD8^+^ or CD4^+^ live singlets. Cells in (B), (D), and (F) are gated on live singlets only.

**Figure S12:**
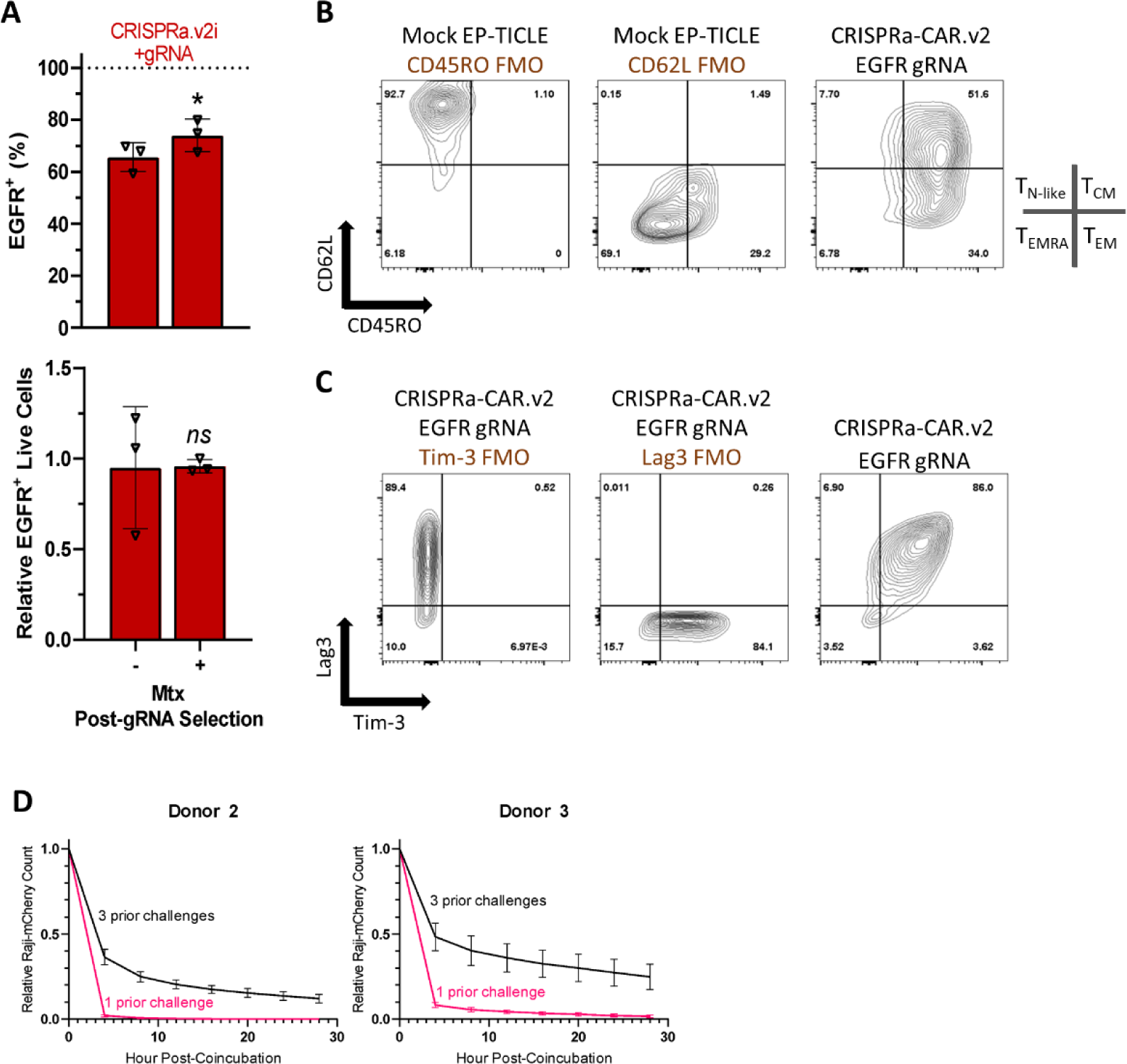
Supporting data for figure 3. (A) Measured EGFR positivity (*top graph*) and total EGFR^+^ cell yields (*bottom graph*) of CRISPRa-CAR.v2i + gRNA-LV T cell as a function of subsequent MTX incubation for 14 days. (B) and (C) Select two-dimensional flow plots measuring expression of indicated antigens (CD45RO, CD62L, Tim-3, Lag3) on CRISPRa-CAR.v2i + gRNA-LV cells quantified in Figure 3 or FMO control. (D) Raji-mCherry CRISPRa-CAR.v2i + gRNA-LV T cell cytotoxicity as a function of prior Raji challenges. Cells in (A), (B), and (C) used 1:1 mixtures of CD8^+^ or CD4^+^ T cells as EP-TICLE input. Cells in (D) used CD8^+^ T cells as EP-TICLE input. All transductions were performed at Day 17 of EP-TICLE manufacturing and assessed four days later via flow cytometry. Cells in (A), (B), and (C) are gated on mScarlet-dCas9^+^CD8^+^ or CD4^+^ live singlets. Data in (A) was evaluated by a paired student t-test. * p < 0.05. ns = not statistically-significant.

**Figure S13:**
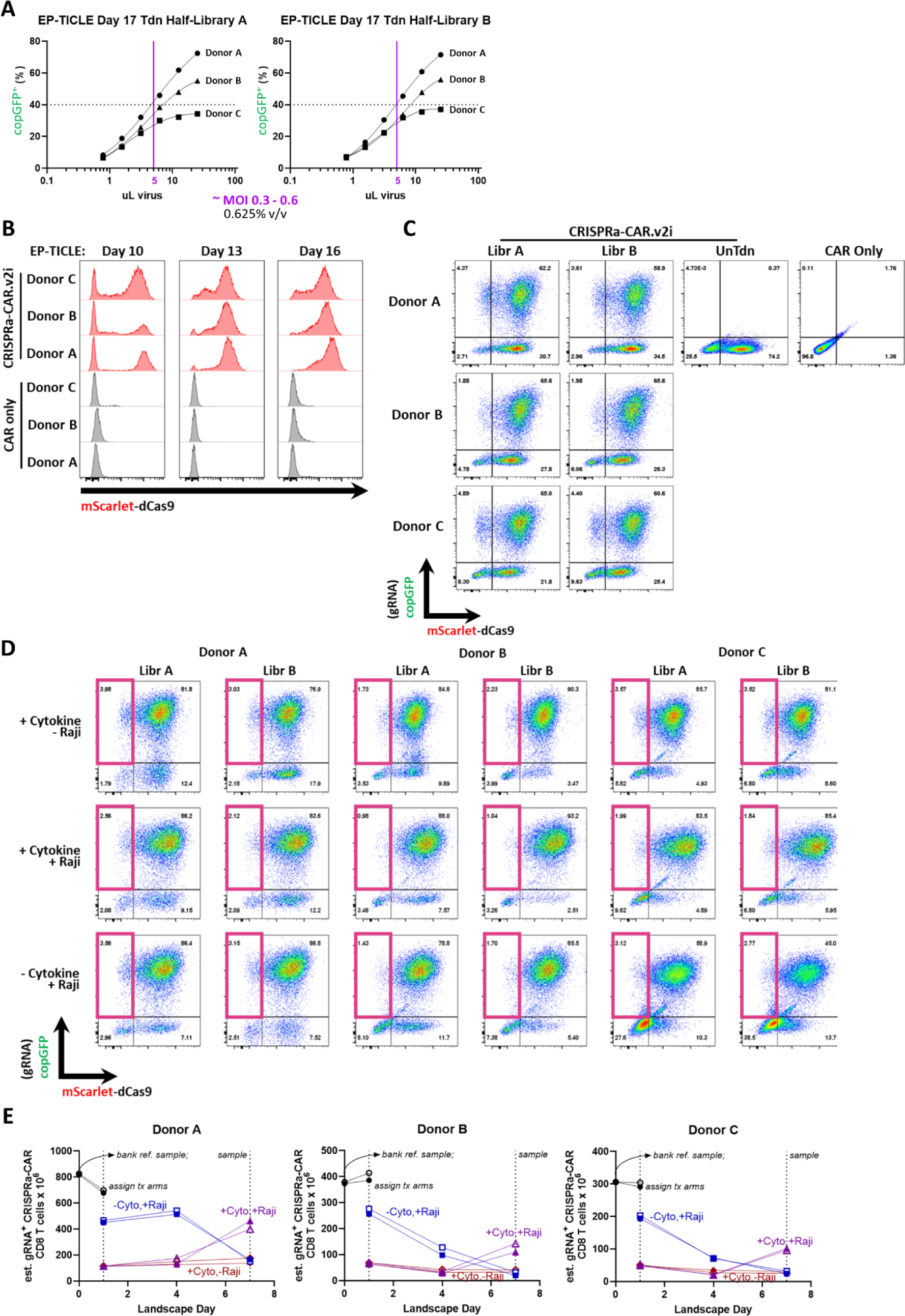
Supporting data for figure 4. (A) Titration of Jensen-Calabrese library lentivirus on Day 17 of EP-TICLE CRISPRa-CAR T cells, separated by indicated donor. (B) mScarlet-dCas9 expression during EP-TICLE manufacturing of CRISPRa-CAR T cells preceding genome-scale screening. (C) mScarlet-dCas9 and copGFP expression at Day 1 of landscape initiation. (D) mScarlet-dCas9 and copGFP (gRNA) expression of CRISPRa-CAR T cell cultures assigned to indicated treatment arms at Day 4 following landscape initiation. (E) Kinetics of CRISPRa-CAR T cell absolute expansion in indicated donor as a function of treatment arm. Cells in (B), (C), and (E) are gated on live singlets. Framed pink quadrant in (D) highlights persistent low percentage of copGFP^+^mScarlet-dCas9^−^ cells.

**Figure S14:**
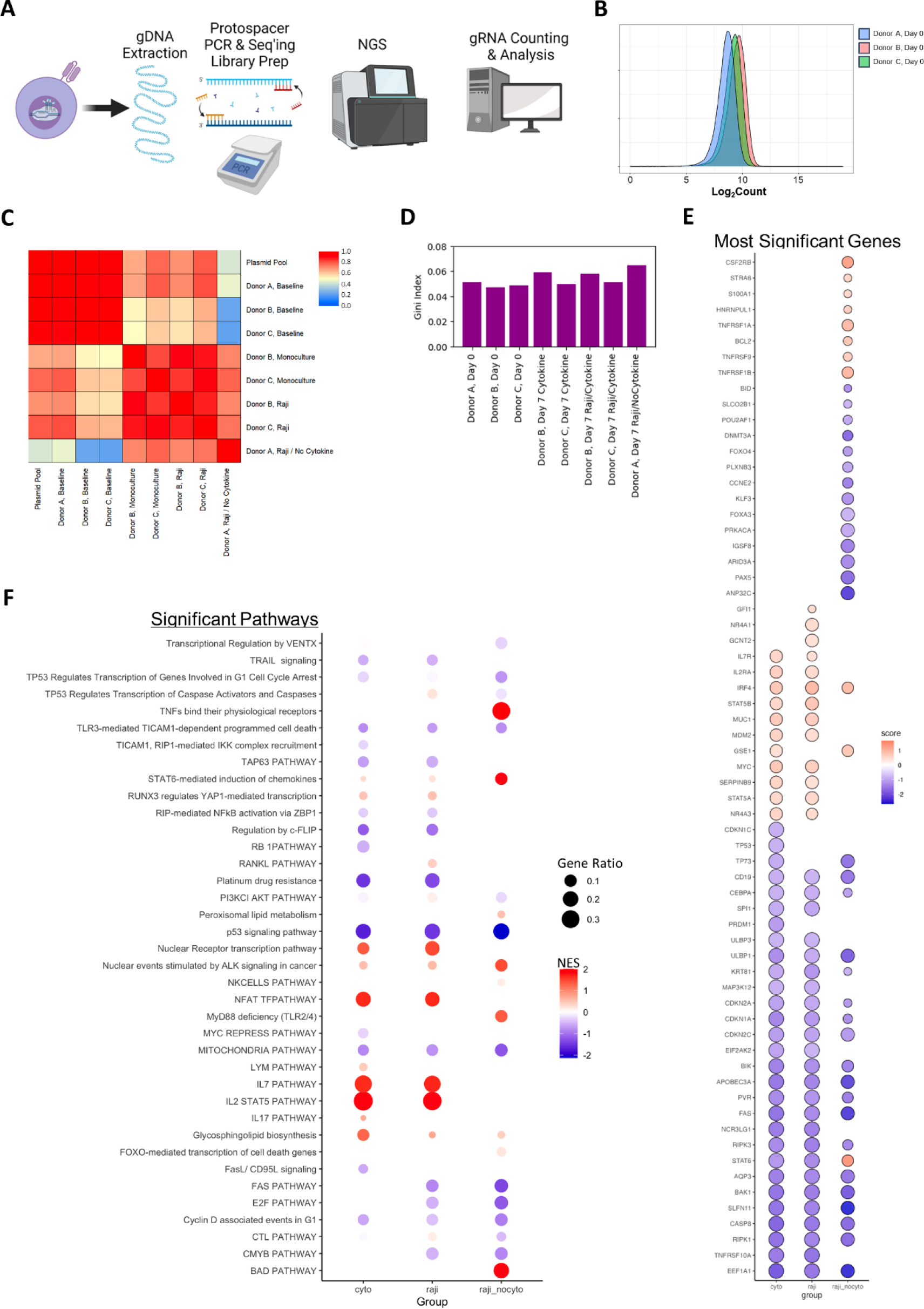
Extended data for figure 4. (A) Graphic illustrating CRISPRa-CAR T cell genome-scale screen sample recovery and subsequent processing. (B) Log_2_ counts for Donors A, B, and C at Day 0 of the genome-scale screen. (C) Pairwise Pearson correlation matrices for indicated samples. (D) Gigi coefficient specific to each indicated sample. (E) Gene-targeting gRNAs significantly enriched or depleted across treatment arms. (F) Gene pathways with significantly enriched or depleted gRNAs across treatment arms.

## Supplemental Materials

Table S1: sequence origins

Table S2: human introns

Table S3: gRNA sequences

Table S4: NGS indexing primers

Table S5: Jensen-Calabrese plasmid pool libraries

Table S6: LV production reagents

Table S7: general reagents

Table S8: flow antibodies

Table S9: ddPCR primer-probes

Table S10: results of genome-wide CRISPRa-CAR *in vitro* persistence, cytokine only (two donors)

Table S11: results of genome-wide CRISPRa-CAR *in vitro* persistence, Raji co-culture with cytokine (two donors)

Table S12: results of genome-wide CRISPRa-CAR *in vitro* persistence, Raji co-culture without cytokine (single donor)

Table S13: GOBP analysis, bottom 200 gene-specific gRNA set pathways, cytokine only

Table S14: Comprehensive significantly-enriched pathways analysis, all treatment arms

## Supplementary Information

File 1: Supplementary Results

File 2: Supplementary Tables S1 through S14

## Author Contributions

BC and MCVJ conceived of the project.

BC, NM, and JM-R cloned constructs used in these studies.

BC, JMR, CPS, LF, AML, and ML isolated T cells used for all studies.

BC, NM, JM-R, and CPS manufactured, manipulated, and characterized T cells.

BC, NM, and AV processed gDNA and prepared sequencing libraries for amplicon-seq.

TC and BC developed ddPCR assays.

CHS, AV, and BC performed computational analyses of sequencing data.

LF, AML, ML, and BC produced and titered all lentivirus used in these studies.

MF provided guidance and critical suggestions.

MCVJ and SKO supervised the project.

BC drafted the initial manuscript.

BC, MCVJ, SKO, TC, CHS, AV, and FL edited the draft manuscript.

All authors reviewed and approved of the final manuscript.

## Acknowledgements

We thank Jacob Appelbaum for valuable conversations related to interrogation of CRISPRa durability; Rachael Logan, Taylor Ishida, and Andrew Gramer for technical assistance with cell handling and other assay development; Samantha Steurer for assistance manufacturing large-scale lentivirus for the genome-scale screen; Marlena Urvater and Pittra Jaenprajak for assistance in early CRISPRa-CAR manufacturing development; Mariliis Ott and Ricardo Miranda for early assay development of the EP-TICLE process; Joshua Gustafson for facilitating the prior collaborations and permitting access to invaluable shared resources; and Els Verhoeyen for kindly providing BaEVRless and H/F plasmids used in this study.

## Data and Materials Availability

Raw sequencing data critical to the findings of this study will be deposited at the National Center for Biotechnology Information (NCBI) Gene Expression Omnibus. Currently additional sequencing studies are pending. Analyzed data are available in the main text of this study or in the supplied supplementary materials.

