## Supplemental Results for "Genome-wide CRISPRa screens nominate modulators of CAR T cell survival within distinct tumor cytokine milieus"

Prior to implementation in primary T cells, we first profiled and refined established CRISPRa CDSs for our purpose using the Jurkat immortalized T cell line. It has been suggested that CRISPRa platforms exhibit lower average levels of gene overexpression relative to more conventional open reading frame (ORF) transgene delivery;^1^ thus, to maximize our screen sensitivity, we engaged in several rounds of parallel and serial optimization. Following this process, we selected a CRISPRa vector featuring an mScarlet-dCas9 fusion (**Fig S1**) separated by a c-Myc nuclear localization signal (NLS) (**Fig S2**). Also incorporated into our final vector was a modified second-generation synergistic activation mediator (SAM) system design,^2^ exchanging a pair of MS2-PP7 hybrid aptamers within the gRNA stems for a pair of pure PP7 aptamers^3^ (**Fig S3**). Lastly, unlike conventional SAM CRISPRa designs, we chose to omit any dCas9-tethered effector due to its ambiguous benefit both in prior studies^3,4^ and in our hands (**Fig S1**).

To deploy these refashioned tools to primary T cells, we first evaluated the feasibility of lentiviral delivery. CRISPRa and CAR transgenes are large DNA payloads, together spanning more than 10 kilobases in total length, preventing their efficient packaging into a single lentiviral viron.^5,6^ Therefore, and similar to previous studies,^7^ we distributed our transgenes across lentiviral pairs (**Fig S4a**). To manufacture experimental (CRISPRa-CAR_gRNA_) or reference (mScarlet-CAR_gRNA_) populations, activated primary T cells were transduced with lentivirus in line with conventional cell manufacturing protocols (**Fig S4c**). Following gene delivery, cells were expanded and efficiently selected (87 – 98%) for CAR positivity. As expected, reference cell populations receiving an mScarlet control lentivirus (mScarlet-LV) achieved moderate levels of mScarlet positivity (55 - 62%); however, in line with prior reports, cells transduced with dCas9-LV registered very low frequencies of dCas9^+^ cells (3 - 6%), despite being transduced at an identical multiplicity of infection (MOI) (= 1) (**Fig S4d**).^8^

Later, to circumvent premature transgene deactivation, we conceived of a CRISPRa Tet-Off design that repressed CRISPRa transcription throughout manufacturing via the supplementation of the Tet repressor, doxycycline (Dox). After successfully piloting this system in Jurkat T cells (**Fig S7a, b**), we transferred our design to CAR-containing constructs, and deployed them to primary T cells.^9^ Disappointingly, primary CRISPRa-CAR T cells featuring this genetic circuit failed to demonstrate rescued CRISPRa expression upon Dox removal (**Fig S7a, d**).

Finally, to maximize transduction efficiency of gRNA-LV, we set about building a simplified and efficient gRNA transfer vector. Existing public CRISPRa gRNA libraries routinely bundle the CRISPRa activator (e.g. PPH) at a substantial cost to lentiviral titer. Having already delivered such components during primary manufacturing, we replaced redundant sequences in the transfer vector with DNA encoding a super-bright green fluorescent protein (copGFP). We then tested varying the positioning of our gRNA cassette, and ultimately selected the PP7 gRNA variant present in our all-in-one CRISPRa-CAR.v1_gRNA_ donor plasmid (Libr.1) due to its superior viral titer and CRISPR activity (**Fig S10a - e**). We also characterized alternative gRNA-LV formats capable of oligo(dT) capture and downstream scRNA-seq processing that are distinct from previous reports;^16^ however, these high content gRNA forms incurred costs to CRISPRa potency (Libr.3) or viral titer (Libr.7), and thus were not adopted for large-scale screening (**Fig S10a - e**).
